# Energy status-promoted growth and development of Arabidopsis require copper deficiency response transcriptional regulator SPL7

**DOI:** 10.1101/2021.09.17.460807

**Authors:** Anna Schulten, Björn Pietzenuk, Julia Quintana, Marcus Krause, Regina Feil, Maida Romera-Branchat, Vanessa Wahl, Edouard Severing, George Coupland, Ute Krämer

## Abstract

Copper (Cu) is a cofactor of around 300 Arabidopsis proteins including photosynthetic and mitochondrial electron transfer chain enzymes critical for ATP production and carbon fixation. Plant acclimation to Cu deficiency requires the transcription factor SQUAMOSA PROMOTER-BINDING PROTEIN-LIKE7 (SPL7). We report that in the wild type and in the *spl7-1* mutant, respiratory electron flux *via* Cu-dependent cytochrome *c* oxidase remained unaffected under both normal and low-Cu cultivation conditions. Contrary to the wild type, supplementing Cu-deficient media with exogenous sugar failed to stimulate growth of *spl7*-*1*. The *spl7*-*1* mutant accumulated carbohydrates including the signaling sugar trehalose 6-phosphate, as well as ATP and NADH, also under normal Cu supply and without sugar supplementation. Late flowering of *spl7-1* was in agreement with its attenuated sugar responsiveness. Functional TOR and SnRK1 kinase signaling in *spl7*-*1* suggested against fundamental defects in these energy-signaling hubs. Sequencing of chromatin immunoprecipitates combined with transcriptomics identified direct targets of SPL7-mediated positive regulation, including *FE SUPEROXIDE DISMUTASE1* (*FSD1*), *COPPER-DEFICIENCY-INDUCED TRANSCRIPTION FACTOR1* (*CITF1*) and uncharacterized *bHLH23* (*CITF2*), as well as an enriched upstream GTACTRC motif. In summary, transducing energy availability into growth and reproductive development requires the function of SPL7. Our results could help to increase crop yields, especially on Cu-deficient soils.

## Introduction

As sessile organisms, plants depend on acquiring mineral nutrients from the soil solution through their roots. Several class B and borderline elements, which are often addressed as “transition metals” or simply “metals”, for example iron (Fe), zinc (Zn), and copper (Cu), are micronutrients required for the functions of numerous metalloproteins. The plant metal homeostasis network operates to fulfill the demands of the metalloproteomes of different organs, tissues and cell types, as well as to counteract local accumulation of a toxic excess of any essential or chemically similar non-essential metal (Clemens, 2001; Krämer and Clemens, 2005). Many of the network components mediating metal acquisition, distribution, utilization and storage in plants were functionally characterized. We know less about regulatory components, for example metal sensors, signal transduction pathways and proteins activating acclimation responses. Among the latter, several identified transcription factors mediate transcriptional responses to Fe, Zn or Cu deficiencies in Arabidopsis (Colangelo and Guerinot, 2004; Wang et al., 2007; Yamasaki et al., 2009; Assunção et al., 2010; Bernal et al., 2012; Li et al., 2016; Liang et al., 2017; Yan et al., 2017).

Of all essential metals, Cu cations possess the highest ligand-binding affinities (Fraústo da Silva and Williams, 2001). As a result, Cu cofactors carry out challenging ligand-binding tasks, for example the interaction with gas molecules. In addition, Cu readily catalyzes single-electron transfer reactions involving the Cu^2+^/Cu^+^ redox couple. Probably as a result of these potent chemical properties, cellular Cu homeostasis is exceptionally tight (Rae et al., 1999; Robinson and Winge, 2010; Foster et al., 2014). In plants, the largest Cu quota are required in chloroplasts where Cu acts as a cofactor in PLASTOCYANIN (PC), the soluble electron carrier between the cytochrome *b_6_f* complex and photosystem I in photosynthetic electron transfer and one of the most abundant proteins in the thylakoid lumen (Redinbo et al., 1994; Schubert et al., 2002). The association of PC with its Cu cofactor occurs post-translationally in the thylakoid lumen and involves the delivery of Cu by PLASTID CHAPERONE1 (PCH1) to the Cu-transporting P-TYPE ATPASE OF ARABIDOPSIS1 (PAA1) of the inner chloroplast envelope membrane, and subsequently Cu transport by PAA2 across the thylakoid membrane (Shikanai et al., 2003; Abdel-Ghany et al., 2005; Blaby-Haas et al., 2014). Cu is also a cofactor in cytochrome *c* oxidase (COX or mitochondrial complex IV), which transfers electrons to oxygen as the terminal electron acceptor of the respiratory electron transport chain in the mitochondria (mETC) (Kadenbach et al., 2000). Cu delivery to its binding sites in the COX1 and COX2 subunits involves the Cu chaperone COX17 and HOMOLOG OF CU CHAPERONE SCO1 (HCC1), which were identified based on their homology to yeast proteins (Attallah et al., 2011; Garcia et al., 2016). The presence of Cu-dependent enzymes at the two key cellular sites of ATP production emphasizes the central relevance of Cu in plant energy metabolism. Whether plants integrate Cu homeostasis and energy metabolism, however, remains unknown.

The transcription factor SQUAMOSA PROMOTER BINDING PROTEIN-LIKE7 (SPL7) operates as a regulator of Cu deficiency-responsive gene expression in *Arabidopsis thaliana* (Yamasaki et al., 2009; Bernal et al., 2012). SPL7 is a member of a transcription factor family characterized by the Squamosa promoter Binding Protein (SBP) domain containing both the nuclear localization signal and the recognition domain for the binding of a GTAC core DNA motif (Cardon et al., 1999; Birkenbihl et al., 2005). The Arabidopsis SPL family comprises 16 proteins constituting subfamily I (SPL1, 7, 12, 14 and 16) and subfamily II, based on size and sequence similarity (Xing et al., 2010). Subfamily II SPL proteins (SPL2 to −6, −8 to −11, −13 and SPL15) have regulatory roles in diverse aspects of plant development including leaf and trichome formation, developmental phase transitions, floral meristem identity and fertility (Unte et al., 2003; Wu and Poethig, 2006; Wang et al., 2008; Yamaguchi et al., 2009; Xing et al., 2010; Yu et al., 2010; Xing et al., 2013; Xu et al., 2016; He et al., 2018). With plant age, a gradual decrease in the abundance of miRNA156, which targets transcripts encoding all subfamily II SPL proteins except SPL8, contributes to increasing levels and cellular activities of these SPL proteins (Wu et al., 2009; Yang et al., 2013; Yu et al., 2013). With the exception of SPL7, the biological functions of subfamily I SPL proteins are less well understood (Stone et al., 2005; Chao et al., 2017; Schulten et al., 2019).

SPL7 is required to enhance the transcription of genes encoding proteins with roles in Cu acquisition, such as root surface Cu(II) chelate reductases (annotated as FERRIC REDUCTASE OXIDASE, FRO) FRO4/FRO5 and transporters of Cu^+^ into the cytosol of the COPT membrane protein family (Bernal et al., 2012). Additionally, SPL7 mediates the microRNA-dependent post-transcriptional downregulation of the levels of transcripts encoding several abundant but non-essential Cu metalloproteins. This aspect of the Arabidopsis Cu deficiency response is thought to reflect an economization strategy that prioritizes the allocation of Cu to essential cuproproteins like PC (Abdel-Ghany and Pilon, 2008; Yamasaki et al., 2009). For example, under Cu deficiency the abundant CuZn superoxide dismutase-encoding *CSD2* and *CSD1* transcripts are targeted by miR398 to replace these CSDs by Fe SUPEROXIDE DISMUTASE1 (FSD1) in an SPL7-dependent manner (Yamasaki et al., 2007; Abdel-Ghany and Pilon, 2008; Yamasaki et al., 2009). The principle of Cu economization was first discovered in the green alga *Chlamydomonas reinhardtii* involving SBP domain-containing CU RESPONSE REGULATOR1 (CRR1) (Quinn and Merchant, 1995; Kropat et al., 2005). ChIP-seq using transgenic *A. thaliana* expressing FLAG-tagged SPL7 under the control of the CaMV 35S promoter identified 1,266 genes associated with SPL7 binding sites under Cu deficiency (Zhang et al., 2014), far more than the 188 transcripts identified as up- or downregulated in an *SPL7*-dependent manner (Bernal et al., 2012).

Here we examine the hypothesis of an integration between Cu homeostasis and plant energy metabolism through *SPL7*. The Arabidopsis *spl7*-*1* mutant, which lacks a broad range of Cu deficiency responses, was unresponsive to growth stimulation by sucrose under Cu deficiency, different from the wild type. An accumulation of sugars in *spl7*-*1*, including also the signaling sugar trehalose 6-phosphate (T6P), suggested that the mutant is impaired in sugar utilization. Indicating against simple biochemical defects in *spl7*-*1*, respiratory electron flux, NADH and ATP levels were normal or even elevated in the mutant. Activities of TOR and SnRK1 kinases were sugar-responsive and in agreement with elevated sugar levels in *spl7-1*, suggesting that these major energy signaling pathways are generally functional in the mutant. The *spl7*-*1* mutant bolted substantially later than the wild type. We sequenced chromatin immunoprecipitates from *A. thaliana spl7-1* producing HA-tagged SPL7 under the control of the native promoter and terminator, and we conducted comparative transcriptomics in *spl7*-*1* and the wild type. In combination, these data suggest a direct activation of transcription of bHLH transcription factor-encoding genes by SPL7 and delineate candidate processes and genes for roles in *SPL7*-dependent sugar signaling. We report a predominantly promoter-localized GTACTRC motif as enriched among SPL7-bound genes showing *SPL7*-dependent increase in transcript levels under Cu deficiency, while other enriched motifs are likely to reflect additional and complex roles of SPL7. In summary, maintaining a balance between energy availability, growth and development requires *SPL7* function, especially under Cu-deficient growth conditions.

## Results

### The *spl7*-*1* mutant is impaired in directing sugars into growth processes and accumulates sugars

To test for interactions between Cu homeostasis and sugar metabolism, we transferred 7-d-old wild-type and *spl7-1* mutant seedlings onto agar-solidified media differing in Cu and sucrose contents (Supplemental Figure 1A and E, compare B-D and F)(Bernal et al., 2012; Marschner and Marschner, 2012). In the wild type, we observed a sucrose-dependent stimulation of biomass production by up to 60% irrespective of Cu supply, but this sugar response was absent in *spl7-1* cultivated in low Cu (Figure 1A-B). Also on moderate sucrose concentrations between 0.1 and 3% in low-Cu medium, biomass of *spl7*-*1* did not increase significantly, in contrast to the wild type (Supplemental Figure 2). Anthocyanin production is a well-known response of plants to high internal levels of sucrose (Larronde et al., 1998; Weiss, 2000; Teng et al., 2005; Solfanelli et al., 2006). The *spl7*-*1* and the *spl7-2* mutants both accumulated higher anthocyanin levels than the wild type and a complemented line (Bernal et al., 2012), with the highest anthocyanin levels in *spl7* mutants cultivated under low Cu conditions (Figure 1A and C, Supplemental Figure 3A). The maintenance of wild type-like anthocyanin levels in *ran1-1* (Woeste and Kieber, 2000) indicated against an ethylene-signaling defect as a cause of this *spl7* mutant phenotype. Shoots of *spl7*-*1* cultivated in low Cu accumulated about 2-fold higher sugar levels than wild-type shoots without exogenous sucrose supply and 17- and 20-fold higher sucrose and glucose levels under high-sucrose cultivation conditions (Figure 1D-E, Supplemental Figure 3B-C for an independent experiment). Sugar levels were elevated in the *spl7-1* mutant even under Cu-sufficient control conditions, despite a fresh biomass comparable to the wild type (Figure 1A-B and D-E, Supplemental Figure 3B-C).

**Figure 1.**
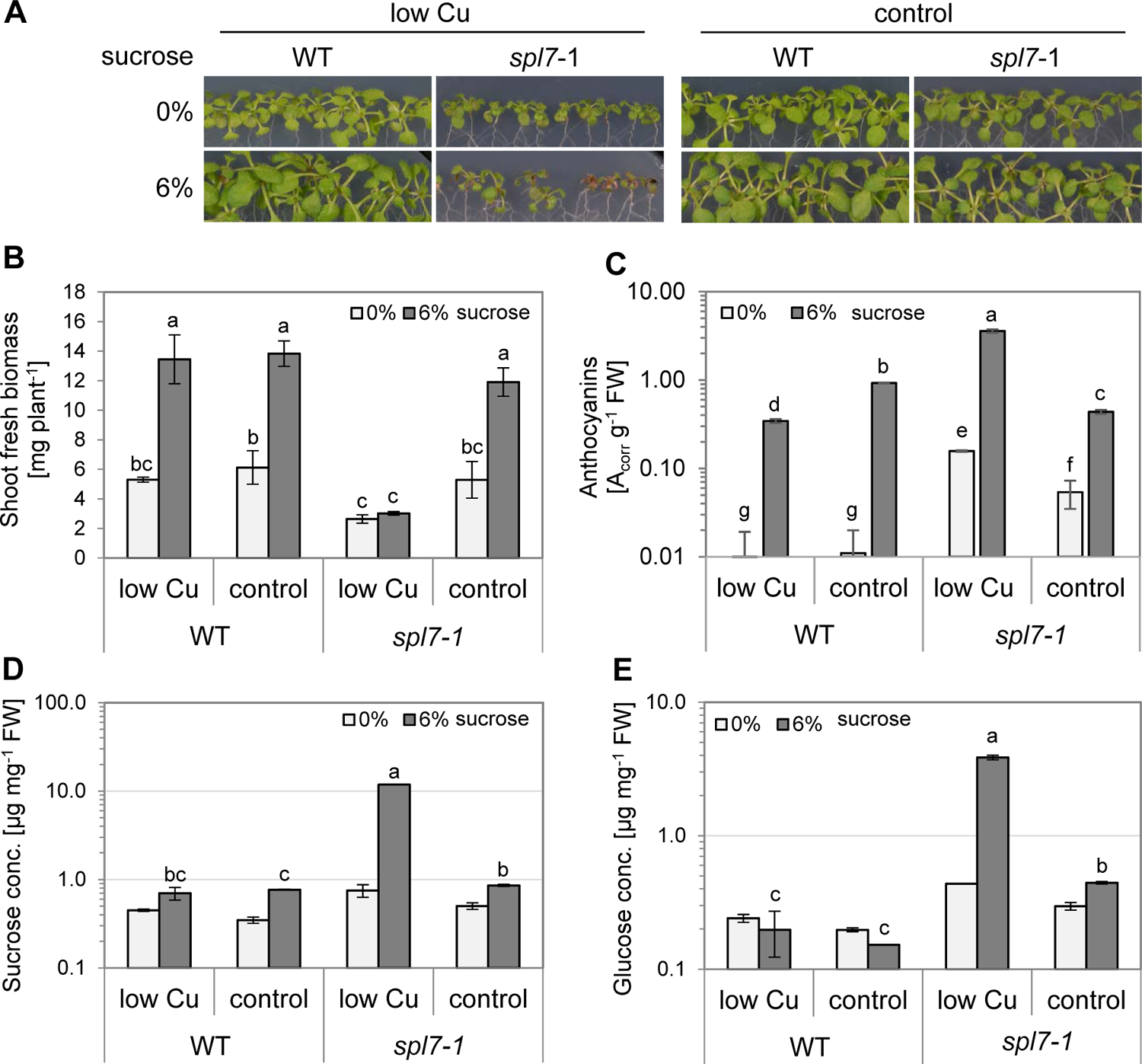
The *spl7-1* mutant accumulates elevated levels of sugars instead of directing them into growth. (A) Photographs of 21-day-old wild-type (WT) and *spl7-1* mutant seedlings cultivated for 14 d in vertically oriented glass petri dishes on low-Cu (0.05 µM CuSO_4_) or control (0.5 µM CuSO_4_) EDTA-washed agar-solidified media supplemented with 0% or 6% (w/v) sucrose. (B) Shoot fresh biomass of seedlings (see A). Bars represent arithmetic means ± SD (*n* = 3 pools of seedlings, with each pool from one replicate plate). (C) Anthocyanin levels in shoots (see A). Bars represent arithmetic means ± SD (*n* = 3 technical replicates for one pool). (**D**, **E**) Sucrose (D) and glucose (E) concentrations in shoots (see A). Bars represent arithmetic means ± SD (*n* = 2 and 3 technical replicates for 0% and 6% sucrose samples, respectively). Data from an independent experiment (C-E) are shown in Supplemental Figure 3. Different characters denote statistically significant differences between means based on ANOVA (Tukey’s HSD; *p* < 0.05, B; *t*-tests with FDR adjustment, *q*-value < 0.05, C-E, wherever *n* > 2). n.d.: not detectable.

We also observed *SPL7*-independent ionomic alterations in high-sucrose grown seedlings, namely generally decreased shoot Fe concentrations and decreased shoot Cu levels only under low-Cu conditions (Supplemental Figure 1A and B, compare with C-F). We confirmed sucrose responses of transcript levels of known *SPL7*-dependently regulated genes in the wild type, which were reported previously on media containing identical sucrose concentrations (Ren and Tang, 2012)(Supplemental Figure 4). These sucrose responses of transcript levels were generally of low magnitude, and they remained detectable in the *spl7-1* mutant.

Pursuing our focus on *SPL7*-dependent phenotypes in this present study, we sought to identify the causes of the defects in sugar utilization in *spl7-1*. In plants, part of the fixed carbon accumulates as starch during the day and is remobilized during the night to avoid carbon depletion and allow continued growth, which is why several mutants with defects in starch synthesis or degradation (e.g. *pgm*) were reported to have reduced growth rates in short days (Caspar et al., 1985; Gibon et al., 2004). At the end of the day, starch concentrations in the *spl7*-*1* mutant paralleled sugar levels, and they were similar or higher (0% sucrose), or far higher (6% sucrose), than in the wild type, demonstrating that the starch biosynthesis pathways are generally functional in the mutant (Supplemental Figure 5A). At the end of the night, starch was depleted or strongly reduced throughout, except in *spl7*-*1* cultivated under low-Cu-high-sucrose conditions (Supplemental Figure 5B). We view this as a consequence of the very high availability of soluble sugars (see Figure 1D and E, Supplemental Figure 3B and C) and starch in the mutant under this condition. We conclude that both the synthesis and mobilization of starch are generally functional in *spl7-1*.

Taken together, these results suggest that *spl7*-*1* is generally impaired in the utilization of sugars and that low-Cu growth conditions exacerbate this further. This could reflect a direct function of SPL7, or it could result indirectly as a symptom from the loss in *spl7-1* of Cu-dependent biochemical functions required for sugar utilization, given the Cu homeostasis defects in this mutant and its apparent susceptibility to physiological Cu limitation.

### Increased respiration rates and altered profile of respiration-related metabolites in *spl7*-*1*

Mitochondrial respiration connects the breakdown of carbohydrates with the production of ATP and of carbon skeletons as biosynthetic precursors (O’Leary et al., 2019). We tested whether defective Cu homeostasis in the *spl7*-*1* mutant results in respiratory restrictions, which could occur through a decrease in Cu-dependent COX activity and feed back to cause insufficient sugar catabolism (Dahan et al., 2014). Notably, total respiration rate was not decreased in *spl7*-*1* and even significantly higher than in the wild type under low Cu (Figure 2A). In the presence of the inhibitor of mitochondrial alternative oxidase (AOX) salicylhydroxamic acid (SHAM) respiration rates did not differ between the *spl7-1* mutant and the wild type (Figure 2A). When AOX activity is inhibited by SHAM, electrons are not redirected into the COX pathway (Bahr and Bonner, 1973; Møller et al., 1988). Consequently, our results suggest that COX (mitochondrial complex IV) was fully functional in *spl7*-*1* even in low-Cu medium.

**Figure 2.**
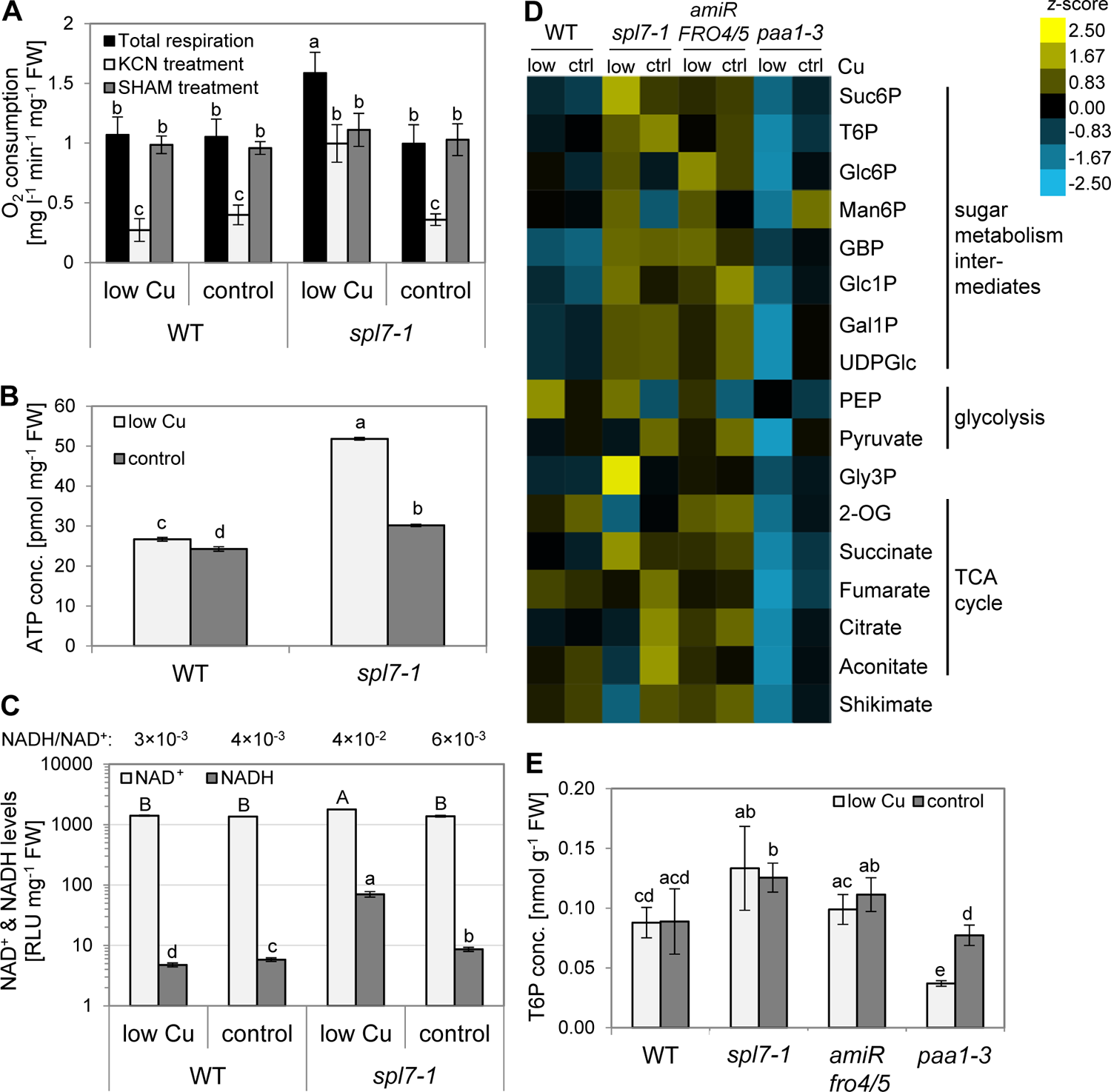
Respiratory activity and quantification of metabolites associated with sugar metabolism and respiration. (**A**) Respiration rates measured as O_2_ consumption in darkness. Data are from leaves of 21-day-old wild-type (WT) and *spl7-*1 seedlings cultivated in vertically oriented glass petri dishes on low-Cu (no CuSO_4_ added) or control (0.5 µM CuSO_4_) EDTA-washed agar-solidified media supplemented with 1% (w/v) sucrose. Complex IV inhibitor KCN (1 mM) or AOX inhibitor SHAM (20 mM) were present during a subset of respiration measurements as indicated. Bars represent arithmetic means ± SD (*n* = 8 and 4 replicate seedling batches for total respiration and for inhibitor treatments, respectively). (**B**, **C**) ATP (B), NAD^+^ and NADH levels (C) in shoots of seedlings cultivated as described for (A). Bars represent arithmetic means ± SD (*n* = 3 technical replicates). (D) Heatmap of metabolites of respiratory and sugar metabolism. Data are from shoots of 21-day-old seedlings (wild-type, *spl7-1*, *35S:amiRFRO4/5*, *paa1-3*) cultivated in vertically-oriented glass petri dishes on low Cu (0.05 µM CuSO_4_) or control (0.5 µM CuSO_4_) EDTA-washed agar-solidified media lacking added sucrose for 14 d. Represented are *z*-scores (*n* = 4 replicate pools per genotype, with one pool per replicate petri plate). (E) T6P concentrations in shoots of seedlings cultivated as described for (D). Bars represent arithmetic means ± SD (*n* = 4, see (D)). Compare Figure 5A. Different characters denote statistically significant differences (*p* < 0.05) between means based on ANOVA (Tukey’s HSD)(A, B) or *t*-tests with FDR adjustment (*q*-value < 0.05) for (C, E), with uppercase and lowercase characters corresponding to different statistical test groups. Data for a second (B, C) and two additional independent experiments (D) are shown in Supplemental Figure 6A-D. RLU: relative light units; Suc6P: sucrose-6-phosphate; T6P: trehalose-6-phosphate; Glc6P: glucose-6-phosphate; Man6P: mannose-6-phosphate, GBP: glucose-1,6-bisphosphate Glc1P: glucose-1-phosphate, Gal1P: galactose-1-phosphate, UDPGlc: uridine diphosphate glucose, PEP: phosphoenolpyruvate, Gly3P: glycerol-3-phosphate, 2-OG: 2-oxoglutarate.

In the presence of KCN, electron flux in the cytochrome *c* pathway is inhibited and consequently redirected to the alternative pathway according to the maximum AOX capacity (Møller et al., 1988). Respiration rates of low-Cu-grown *spl7*-*1* remained more than twice as high compared to the wild type under KCN treatment, indicating a much higher capacity for KCN-insensitive respiration in *spl7*-*1* (Figure 2A). Accordingly, elevated levels of total respiration in *spl7*-*1* cultivated under low-Cu conditions can likely be attributed to a higher AOX activity, in agreement with increased transcript levels of all four genes encoding AOX isoforms in shoots of the *spl7*-2 mutant, particularly of the isoform *AOX1D* (Bernal et al., 2012).

The *spl7-1* mutant contained elevated levels of ATP under control conditions, and more pronouncedly under low-Cu conditions, by comparison to the wild type (Figure 2B, Supplemental Figure 6A). The NADH/NAD^+^ ratio of *spl7-1* was between about 1.5-fold (control Cu) and 9-fold (low Cu) higher than that of the wild type (Figure 2C, Supplemental Figure 6B). These ratios are generally consistent with previous findings that NADH contributes less than about 1% of the cytosolic NAD pool in wild-type plants (Heineke et al., 1991; Shen et al., 2006). Compared to the wild type, the *spl7*-*1* mutant contained higher levels of both NAD^+^ and NADH under low Cu (Figure 2C, Supplemental Figure 6B). Taken together, our observations are consistent with a simple model in which the high availability of sugars in *spl7*-*1* feeds into the production of reductant in the form of NADH. The high levels of NADH in turn fuel respiratory electron flow and ATP production, with normal levels of COX activity and increased AOX activity. AOX partially uncouples NADH oxidation in the mETC from ATP production and can thus function in redox balancing (Vanlerberghe, 2013), but its increased activity in *spl7-1* may be insufficient to prevent the excess of NADH and ATP. This model does not yet provide an explanation for the accumulation of sugars in the *spl7-1* mutant.

We analyzed the levels of metabolites associated with respiratory and sugar metabolism in shoots of 21-d-old seedlings grown in low-Cu and control agar-solidified media lacking added sucrose. In addition to wild-type and *spl7-1* seedlings, we included two previously characterized Cu-deficient Arabidopsis lines, the *paa1-3* mutant (Shikanai et al., 2003; Abdel-Ghany et al., 2005) and a transgenic *35S:amiR-FRO4/5* line (Bernal et al., 2012). In *paa1-3*, severe photosynthetic defects resulting from impaired chloroplastic Cu import and compromised plastocyanin function can be rescued by high levels of exogenous Cu. *35S:amiRFRO4/5* lines are partially impaired in the reduction of Cu(II) to Cu(I) at the cell surface, which is required for high-affinity cellular Cu uptake in low-Cu medium (Bernal et al., 2012).

We observed few and quantitatively minor changes in metabolite concentrations in the wild type under Cu deficiency compared to control conditions, none of which were consistent across independent experiments (Figure 2D, Supplemental Figures 6C-D, Supplemental Table 1). The majority of metabolites were present at higher levels in *spl7*-*1* than in the wild type under both low-Cu and Cu-sufficient cultivation conditions. Compared to the wild type, we observed the consistently largest alterations in *spl7-1* for the levels of glycerol-3-phosphate (Gly3P), 2-oxoglutarate (2-OG), shikimate, sucrose-6-phosphate (Suc6P) and succinate, especially under low Cu. The profiles of these metabolites in *spl7-1* differed from *35S:amiR-FRO4/5*, whereas other sugar- and glycolysis-related metabolites showed similar alterations in both *spl7-1* and *35S:amiR-FRO4/5* compared to the wild type (e.g. glucose-1-phosphate [Glc1P] and glucose-1,6-bisphosphate [GBP]). Different from *spl7-1*, the concentrations of nearly all of the analyzed metabolites were markedly lower in *paa1-3* cultivated in low-Cu media. Chloroplasts of *spl7-1* contain about 20% less Cu than those of the wild type (Zhang et al., 2014), whereas a 58% reduction in chloroplast Cu levels was reported for *paa1-3* (Abdel-Ghany et al., 2005), suggesting a more severe lack of Cu in chloroplasts of *paa1-3*. The opposing trends in metabolite profiles between *spl7-1* and *paa1-3*, together with specific alterations in *spl7-1* when compared to *35S:amiR-FRO4/5*, suggested that the contributions of chloroplast and general physiological Cu deficiency to the metabolite profiles of *spl7*-*1* were minor.

Notably, both of the most strongly altered metabolites, Gly3P (57% increase in *spl7*-*1* compared to wild type in low Cu) and 2-OG (60% decrease in *spl7*-*1*), have roles in the shuttling of reducing agents, for example NADH, across the inner mitochondrial membrane as part of the glycerol phosphate shuttle (Shen et al., 2006) and the malate/aspartate shuttle (Journet et al., 1981). The alterations in the levels of Gly3P and 2-OG in *spl7*-*1* are consistent with the mutant being locked in an overall more reduced state.

In accordance with elevated sugar concentrations in *spl7*-*1* (see Figure 1D, E), the levels of all monitored sugar metabolism intermediates were higher in *spl7*-*1* under low Cu compared to the wild type in at least two out of three independent experiments, with Suc6P levels increased by 76%, for example (Figure 2D, Supplemental Figures 6C-D, Supplemental Table 1). Relative to the wild-type, we observed increased levels of the signaling sugar trehalose 6-phosphate in *spl7-1* (T6P; 56% increase compared to wild type) independent of Cu supply in two out of three independent experiments, and T6P levels broadly followed sugar levels, as is known (Lunn et al., 2006). Consequently, growth defects in *spl7-1* cannot be attributed to a depletion in the signaling sugar T6P (Schluepmann et al., 2003; Figueroa and Lunn, 2016). Together, these results suggest that the primary metabolite profile of *spl7-1* is rather different from that of other Cu-deficient mutants and thus implicate *SPL7*-dependent processes, rather than pleiotropic defects common to Cu-limited plants, in reduced growth despite a high energy status of *spl7-1*.

### Major energy signaling pathways are generally functional in the *spl7-1* mutant

Next, we examined sugar-dependent signaling that is known to connect energy status with growth and development (Baena-Gonzalez and Hanson, 2017). Two central regulators in plant energy signaling are the antagonistically operating kinases Target of Rapamycin (TOR) and Sucrose non-fermenting-Related Kinase (SnRK1). SnRK1 is activated in response to a low energy status, acting to repress energy-consuming processes and growth, and it is inhibited by sugars, likely in the form of sugar phosphates such as T6P (Baena-González et al., 2007; Zhang et al., 2009; Nunes et al., 2013a; Zhai et al., 2018). Transcriptional markers for SnRK1 activity were reported for protoplasts transiently expressing SnRK1 subunits as well as for seedlings after short-term sugar starvation treatments (Baena-González et al., 2007; Nunes et al., 2013a). To test for the functionality of the SnRK1-activating pathway in *spl7*-1, we employed a submerged liquid cultivation system in flasks as described, which allows the rapid exchange of media for 3-h sucrose starvation treatments (Nunes et al., 2013a). In response to sucrose starvation, transcript levels of the SnRK1 marker *Dark Inducible 6* (*DIN6)* were strongly upregulated as expected in both the wild type and *spl7-1*, irrespective of Cu supply (Figure 3A-B). This indicated an ability to activate SnRK1 throughout. However, transcript abundance of *DIN6* was clearly lower in *spl7-1* compared to wild type under low-Cu conditions. Transcript levels of *Expansin 10* (*EXP10)*, which are downregulated by increased SnRK1 activity, decreased in response to sucrose starvation in all samples except for *spl7-1* under low Cu (Figure 3C). Together, these observations suggest that SnRK1 activity reflected sugar levels in *spl7-1*, congruent with our interpretation that the SnRK1 pathway is sugar-responsive and thus generally functional in the mutant.

**Figure 3.**
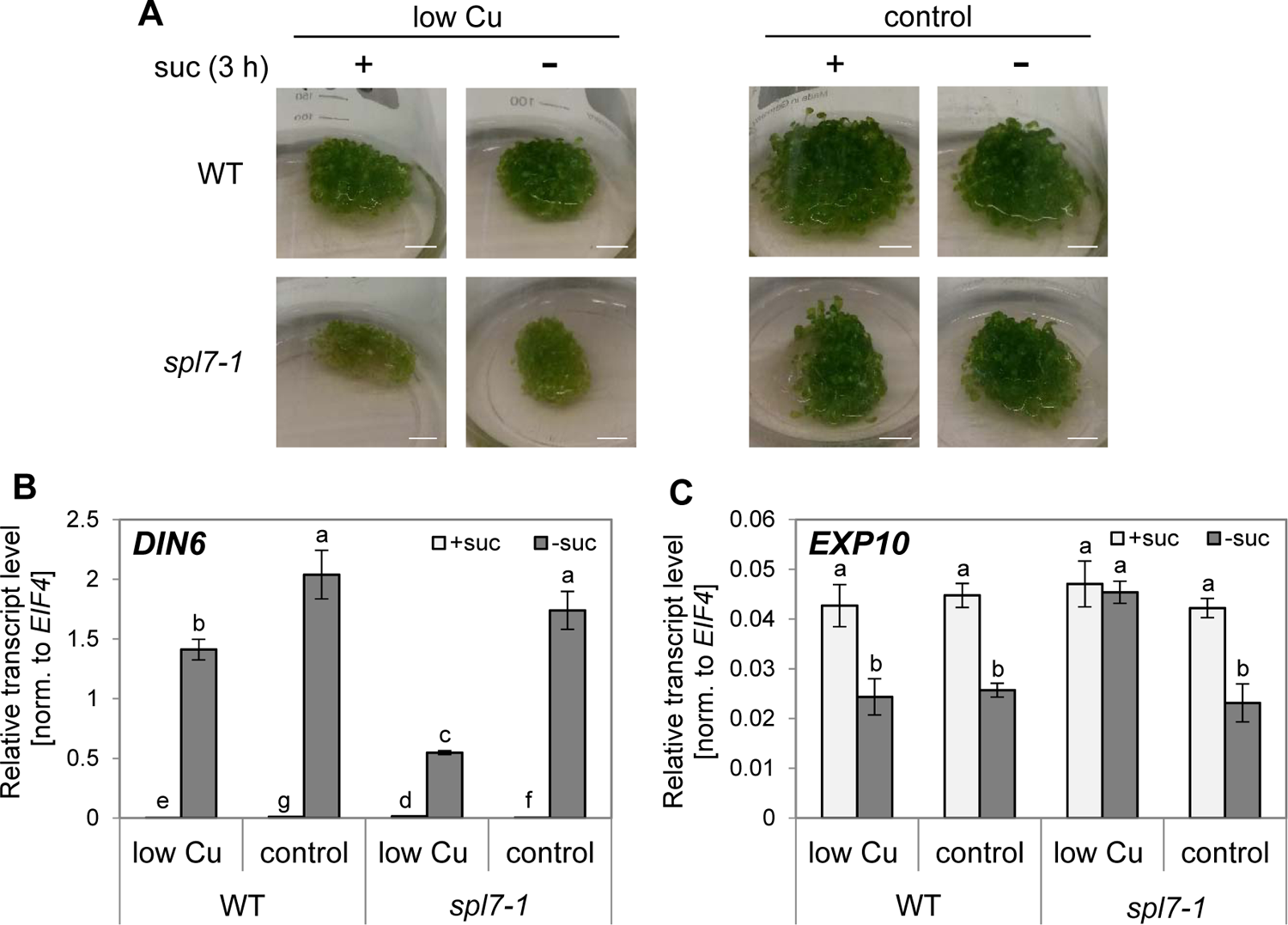
SnRK1 pathway activation in response to sucrose starvation in wild-type and *spl7-1* mutant seedlings. (**A**) Photographs of 15-day-old seedlings cultivated in liquid culture in low-Cu (no CuSO_4_ added) or control (1 µM CuSO_4_) conditions with 0.5% (w/v) sucrose following a 3-h sucrose starvation treatment (0% sucrose). Scale bar, 1 cm. (**B**, **C**) Relative transcript levels of the SnRK1 pathway marker genes *DIN6* (induced by SnRK1 activity) and *EXP10* (repressed by SnRK1 activity) in shoots of seedlings cultivated as described for (A), as quantified by RT-qPCR. Bars represent arithmetic means ± SD (*n* = 3 technical replicates, i.e. independent PCR runs, each with three replicate wells per transcript). Data shown are transcript levels relative to *EIF4* as a constitutively expressed control gene. Different characters denote statistically significant differences (*p* < 0.05) between means based on *t*-tests with FDR adjustment (*q*-value < 0.05)(B) or ANOVA (Tukey’s HSD)(C).

TOR kinase activity is stimulated by sugars, and this promotes growth and development *via* the regulation of processes such as cell proliferation and growth, ribosome biogenesis and protein synthesis (Deprost et al., 2007; Xiong et al., 2013). To assess TOR activity, we analyzed the phosphorylation state of its downstream target S6 kinase (S6K) as a marker in seedlings cultivated in our combined Cu (low Cu and control) and sucrose (0% and 6%) growth conditions, using modification-specific anti-S6K antibodies (Dong et al., 2017). In seedlings grown in 6% sucrose, compared to 0% sucrose, the ratio of phosphorylated relative to non-phosphorylated S6K protein responded no less in *spl7*-1 than it responded in wild-type seedlings, independent of Cu supply (Figure 4, Supplemental Figure 7). This observation is consistent with the known activation of TOR by glucose and sucrose and indicated that TOR-mediated sugar signaling is generally functional in *spl7-1* (Xiong and Sheen, 2012; Xiong et al., 2013; Dobrenel et al., 2016). Following internal sugar levels (see Figure 1D-E), the relative abundance of phosphorylated S6K protein was higher in *spl7-1* than in the wild type upon cultivation in 6% sucrose, and the difference between genotypes was more pronounced and more consistently observed when seedlings were grown in low-Cu medium (Figure 4, Supplemental Figure 7). Taken together, our data implicate a process that acts either independently, or downstream, of the TOR and SnRK1 kinases, in *SPL7*-dependent energy metabolism.

**Figure 4.**
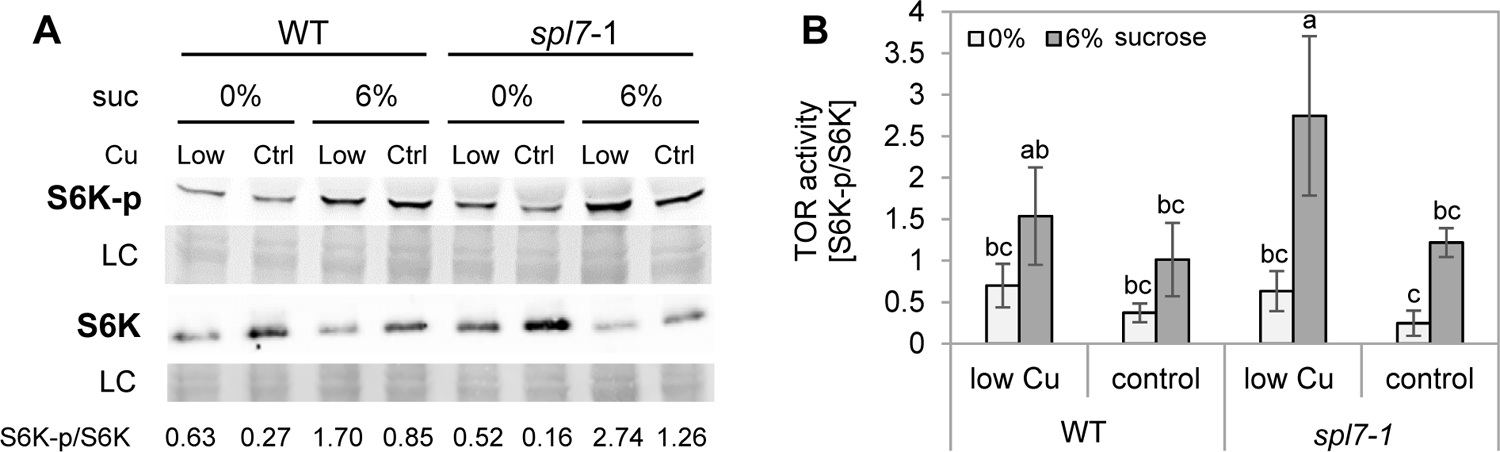
TOR activity in wild-type and *spl7-1* mutant seedlings. (A) Immunodetection of phosphorylated (S6K-p) and total S6K (S6K1 and S6K2) protein as a marker of TOR activity. Data are from shoots of 21-day-old wild-type (WT) and *spl7-1* seedlings cultivated in vertically oriented glass petri dishes on low-Cu (0.05 µM CuSO_4_) or control (0.5 µM CuSO_4_) agar-solidified media supplemented with 0% or 6% (w/v) sucrose for 14 d. Total protein extracts were separated in denaturing polyacrylamide gels and transferred to nitrocellulose membranes. Proteins were visualized on the membrane through Ponceau Red staining as a loading control (LC) prior to immunodetection (S6K-p, S6K apparent sizes 52 kDa). The ratio of S6K-p/S6K band intensities is shown below each lane. Blots from two additional independent experiments are shown in Supplemental Figure 7. (B) Ratios of S6K-p/S6K band intensities for immunoblot images shown in (A) and Supplemental Figure 7. Bars represent arithmetic means ± SD (*n* = 3 replicate blots from independent experiments). Different characters denote statistically significant differences (*p* < 0.05) between means based on ANOVA (Tukey’s HSD).

### Flowering time of *spl7-1*

Sugar signals such as T6P promote growth as well as developmental phase transitions, for example flowering (Schluepmann et al., 2003; Wahl et al., 2013; Yang et al., 2013; Yu et al., 2013; Ponnu et al., 2020). We thus tested for phenotypic changes in *spl7-1* at later developmental stages. For comparison, we included the transgenic *35S:amiRTPS1* line in which T6P levels are reduced as a consequence of the post-transcriptional downregulation of *TREHALOSE 6-PHOSPHATE SYNTHASE1* gene expression, resulting in a strongly delayed flowering time in long days (Wahl et al., 2013). Under our growth conditions, T6P levels were reduced by about 34% in *35S:amiRTPS1* compared to wild type independent of Cu supply (Figure 5A), in agreement with published data (Wahl et al., 2013). In *spl7*-*1* mutant plants cultivated alongside, T6P levels were increased to about 178% of the wild type on low-Cu medium and to 140% of the wild type on control medium.

**Figure 5.**
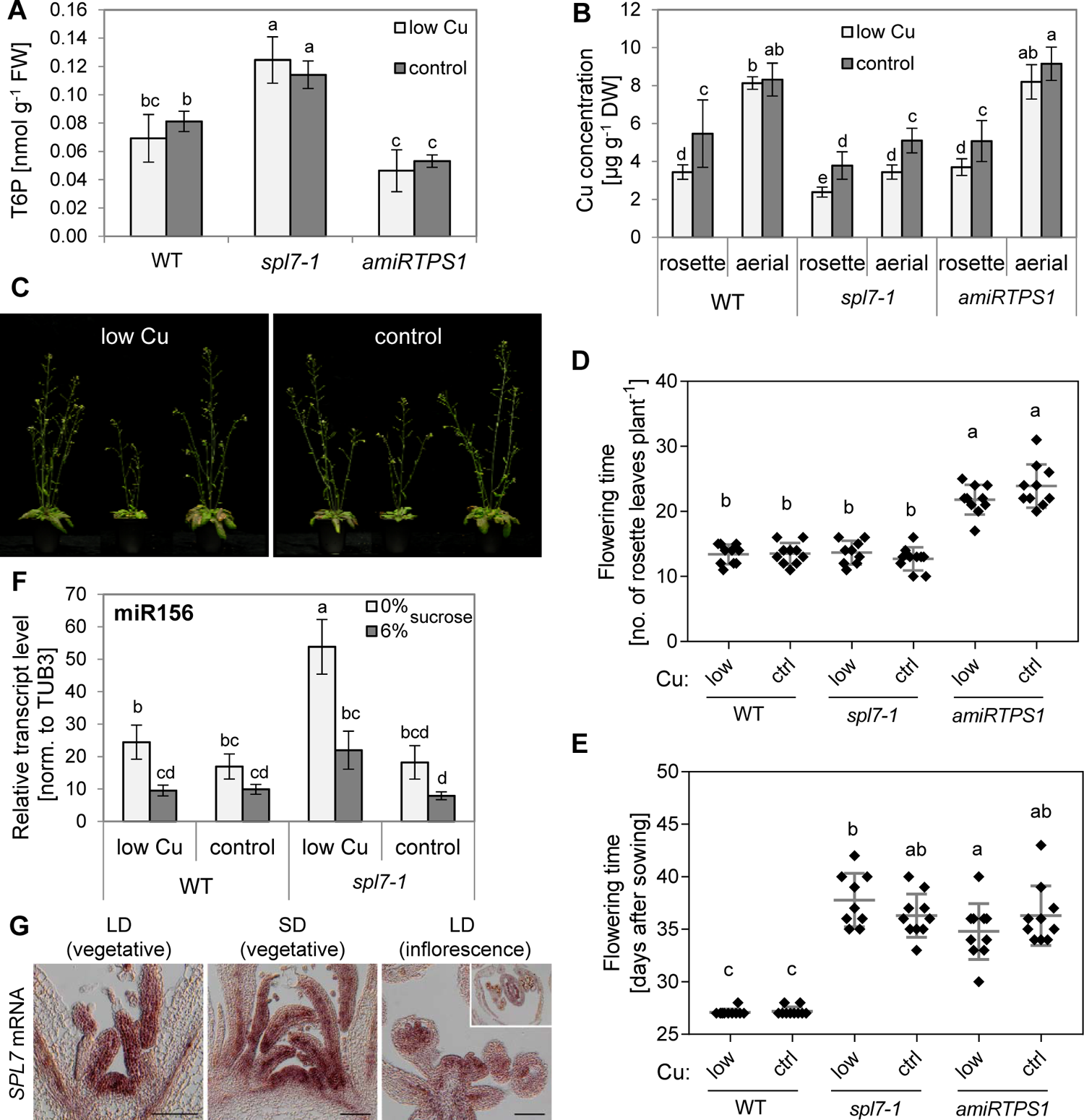
Flowering time of wild-type, *spl7-1* and *3 :am PS1* Ar opsis plants. (A) T6P concentrations. Data are from shoots of 21-day-old see s of wild-type (WT), *spl7-1* and *35S*:*amiRTPS1* cultivated in vertically oriented glass petri dishes on low-Cu (0.05 µM CuSO_4_) or control (0.5 µM CuSO_4_) agar-solidified media without added sucrose for 14 d. Bars represent arithmetic means ± SD (*n* = 4 replicate pools, with one pool per petri plate). Compare Figure 2E (B) Cu concentrations in rosette and aerial tissues of plants cultivated in soil in long days. Plants were watered with equal amounts of tap water without (low Cu) or with 2 mM CuSO_4_ (control) once per week and harvested 10 d after bolting. Bars represent arithmetic means ± SD (*n* = 8 plants per genotype, treatment and tissue). (C) Photographs of wild-type, *spl7-1* and *35S:amiRTPS1* plants 10 d after bolting. Plants were cultivated as described for (B). (**D**, **E**) Flowering time given as the number of rosette leaves (D) and age at bolting time (E) of plants cultivated as described for (B). Plots show arithmetic mean (line) ± SD and all data points (diamonds) (*n* = 8 plants per genotype and treatment). Two independent experiments are shown in Supplemental Figure 8. (F) Relative transcript levels of miR156 in shoots of 21-day-old seedlings. Cultivation was in vertically oriented glass petri dishes on-low Cu (0.05 µM CuSO_4_) or control (0.5 µM CuSO_4_) agar-solidified media supplemented with 0% or 6% (w/v) sucrose for 14 d, with harvest at ZT 3. Bars represent arithmetic means ± SD (*n* = 3 technical replicates, i.e. independent PCR runs, each with three replicate wells per transcript) as calculated based on RT-qPCR. Data shown are representative of two independent experiments. (G) *SPL7* mRNA detection in the shoot apex by *in situ* RNA hybridization. Shown longitudinal sections through apices of wild-type plants cultivated in soil in short (SD) or long-day (LD) conditions. Inset shows a transverse section of a flower. Scale bars, 100 µm. Different characters denote statistically significant differences between means (*t*-tests with FDR adjustment; *q*-value < 0.05)(A, B, D-G). T6P: trehalose-6-phosphate.

Our standard greenhouse soil is Cu-deficient (low Cu), and the defects in reproduction of *spl7-1* are rescued by watering with 2 mM CuSO_4_ (control) once per week (Schulten et al., 2019). With this supplementation, Cu concentrations in rosette leaf and aerial tissues of *spl7-1* were significantly higher than in unamended low-Cu soil (Figure 5B), and the stunted growth of *spl7-1* was partially rescued (Figure 5C), as was reported earlier (Yamasaki et al., 2009; Bernal et al., 2012; Garcia-Molina et al., 2014a; Yan et al., 2017). Cu supplementation led to a significant increase in rosette Cu levels to about 5 µg g^-1^ dry biomass in the wild type and *35S:amiRTPS1*. Cu concentrations in aerial tissues of wild type and *35S:amiRTPS1* were around 8 µg g^-1^ DW, clearly above those in rosette tissues and unaffected by Cu supplementation. By contrast, Cu concentrations in aerial tissues of *spl7-1* were only a little higher than in rosette tissues. This suggested that during the reproductive phase of the life cycle, wild-type plants preferentially allocate Cu into aerial tissues and that this is largely *SPL7*-dependent.

As expected, *35S:amiRTPS1* plants flowered later than the wild type according to both plant age and the number of rosette leaves at bolting (Figure 5D-E, Supplemental Figure 8) (Wahl et al., 2013). Based on the number of leaves at bolting, the *spl7-1* mutant flowered at a similar time or slightly earlier than the wild type (Figure 5D, Supplemental Figure 8A and C). In relation to plant age, bolting time was clearly delayed in *spl7-1* by about 7 days compared to wild type irrespective of Cu supply (Figure 5E, Supplemental Figure 8B and D), and comparable to *35S:amiRTPS1*. Taken together, these results suggest that, in addition to growth, the transition to flowering is partially uncoupled from sugar levels in *spl7-1*. Consistent with a known repression of miR156 by metabolizeable sugars and T6P (Wahl et al., 2013; Yang et al., 2013; Yu et al., 2013; Ponnu et al., 2020), miR156 levels were lower in shoots of 6% sucrose-than in 0% sucrose-grown seedlings of both the wild type and *spl7-1* (Figure 5F). This confirmed that miR156 levels remained responsive to internal sugar levels in *spl7-1* (Yang et al., 2013; Yu et al., 2013). However, despite elevated sugar levels in *spl7-1* compared to the wild type (see Figures 2 and 5A), miR156 levels were higher in *spl7-1* than in the wild type when cultivated under low-Cu conditions (Figure 5F). Thus, the known sugar-dependent decrease in miR156 levels with plant age was attenuated in the mutant in low Cu.

Flowering requires a developmental switch in the apical meristem. Therefore, we investigated whether *SPL7* might be capable of influencing this switch also locally. RNA *in situ* hybridization revealed the presence of *SPL7* transcript in the shoot apical meristem and young leaf primordia of vegetative plants cultivated in both short and long days as well as in reproductive shoot apices (Figure 5G). Several miR156-targets, for example *SPL9*, *SPL13* and *SPL15*, are also expressed in the shoot apical meristem (Wang et al., 2009; Yamaguchi et al., 2014b; Hyun et al., 2016; Xu et al., 2016). The function of *SPL7* in the shoot apical meristem warrants further investigation.

### Genome-wide identification of SPL7 binding sites

We reasoned that expanding our knowledge on direct target genes of SPL7-dependent transcriptional regulation could provide mechanistic insights into how SPL7 affects energy metabolism. For the genome-wide identification of SPL7 DNA binding sites using ChIP-seq, we generated *spl7*-*1* transformants with a genomic construct encoding an SPL7 protein flanked by HA-tags at both the N- and the C-terminus, under control of the native promoter in the *spl7*-*1* background (*promSPL7::HA-SPL7-HA:termSPL7*; short *HA-SPL7-HA*). Of three homozygous lines complementing the *spl7* phenotype under low Cu cultivation conditions, we chose a line in which *SPL7* transcript levels were similar to the wild type (Figure 6A-B, Supplemental Figure 9). *SPL7* transcript levels were unaffected by Cu levels in the medium, as reported previously (Yamasaki et al., 2009; Bernal et al., 2012). Likewise, immunoblots detected the HA-SPL7-HA fusion protein under both low Cu and control conditions (Figure 6C, Supplemental Figure 10), in line with a proposed post-translational mechanism for the regulation of the activity of SPL7 as a transcriptional enhancer, similar to CrCRR1 (Kropat et al., 2005; Sommer et al., 2010). The HA-SPL7-HA protein was visible as a single band running at a size of around 100 kDa, which is slightly higher than the calculated protein size including the HA-tags of ∼92 kDa (Figure 6C, Supplemental Figure 10). Chromatin Immunoprecipitation (ChIP) was conducted on rosette tissues of 21-d-old *HA-SPL7-HA* seedlings cultivated in low-Cu and control conditions. ChIP-qPCR confirmed an enrichment relative to chromatin input for positive control SPL7 target gene *MIR408* (Zhang and Li, 2013) and the expected SPL7 target gene *FSD1* (Zhang et al., 2014) in *HA-SPL7-HA* samples relative to the wild type, in contrast to the negative control gene *ACTIN7* (*ACT7*) (Figure 6D). Next, we identified putative SPL7 binding sites genome-wide based on sequencing of input and ChIP sample pairs. In low-Cu conditions and control Cu conditions, respectively, we identified 758 peaks/2,026 genes and 713 peaks/1,901 genes as SPL7 binding sites/in the vicinity of SPL7 binding sites, supported by a minimum of 2 (of 4) independent experiments (Figure 7A and B, Supplemental Data Sets 1 and 2). About 85% of the genomic segments we identified by ChIP-seq were common to both cultivation conditions (Figure 7B). Around 9% of the genes associated with these genomic segments were among 1,266 genes previously identified based on a single SPL7-ChIP-seq experiment (Zhang et al., 2014)(more than expected by chance; *P* < 10^-15^, hypergeometric test; Supplemental Table 2). Different from the earlier study by Zhang et al. (2014), by far the most frequently observed localization of peaks was 200 to 20 bp upstream of the predicted transcriptional start site (Supplemental Figure 11). Possible explanations for findings differing between the earlier publication and this present study may lie in differing plant cultivation conditions, the earlier expression of FLAG-SPL7 under the control of the CaMV 35S promoter, or the fact that these FLAG-SPL7-ChIP-seq data were apparently based on a single replicate (Zhang et al., 2014).

**Figure 6.**
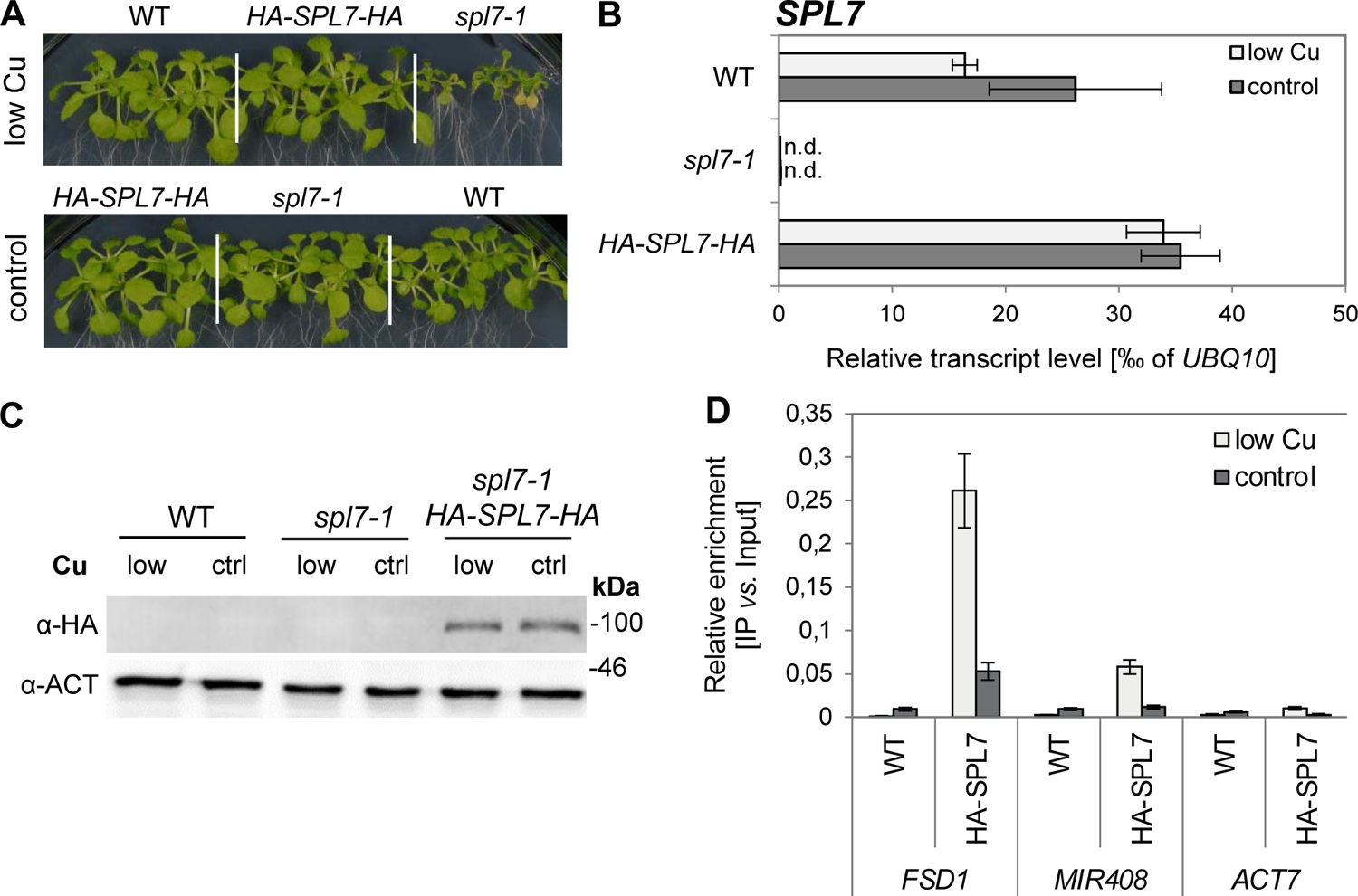
Complementation of the *spl7-1* mutant through the *P_SPL7_::HA-SPL7-HA:t_SPL7_* transgene. (A) Photographs of the wild type (WT), the *spl7-1* mutant and the chosen transgenic homozygous *spl7-1 P_SPL7_::HA-SPL7-HA:t_SPL7_* (*HA-SPL7-HA*) line. Shown are 21-day-old seedlings cultivated in low-Cu (no CuSO_4_ added) or control (0.5 µM CuSO_4_) media supplemented with 1% (w/v) sucrose and solidified with EDTA-washed agar in vertically oriented glass petri dishes. (B) Relative transcript abundance of *SPL7* according to RT-qPCR, in shoots of seedlings cultivated as described for (A). Bars represent arithmetic means ± SD (*n* = 3 technical replicates, i.e. independent PCR runs, each with three replicate wells per transcript). Data from two additional transgenic lines are shown in Supplemental Figure 9. (C) Immunoblot detection of HA-SPL7-HA protein (expected at ∼90 kDa) in shoots of wild-type, *spl7-1* and *HA-SPL7-HA* seedlings cultivated as described for (A). PVDF membranes were stripped and reprobed with α-ACTIN (ACT) antibody (shown as a loading control). The full image and results from an independent experiment are shown in Supplemental Figure 10. (D) Validation of ChIP using ChIP-qPCR of previously implicated direct target genes of SPL7. Bargraph shows relative DNA enrichment of the promoter regions of putative target *FSD1, MIR408* as a positive control gene, as well as *ACT7* (negative control gene), quantified by ChIP qPCR on immunoprecipitation (IP) and input samples. Chromatin was isolated from shoot tissues of *spl7-1 P_SPL7_::HA-SPL7-HA:t_SPL7_* (HA-SPL7) and wild-type seedlings (WT, negative control), cultivated as described for (A). Input samples represent aliquots taken after chromatin shearing and before the addition of α-HA for the IP.

**Figure 7.**
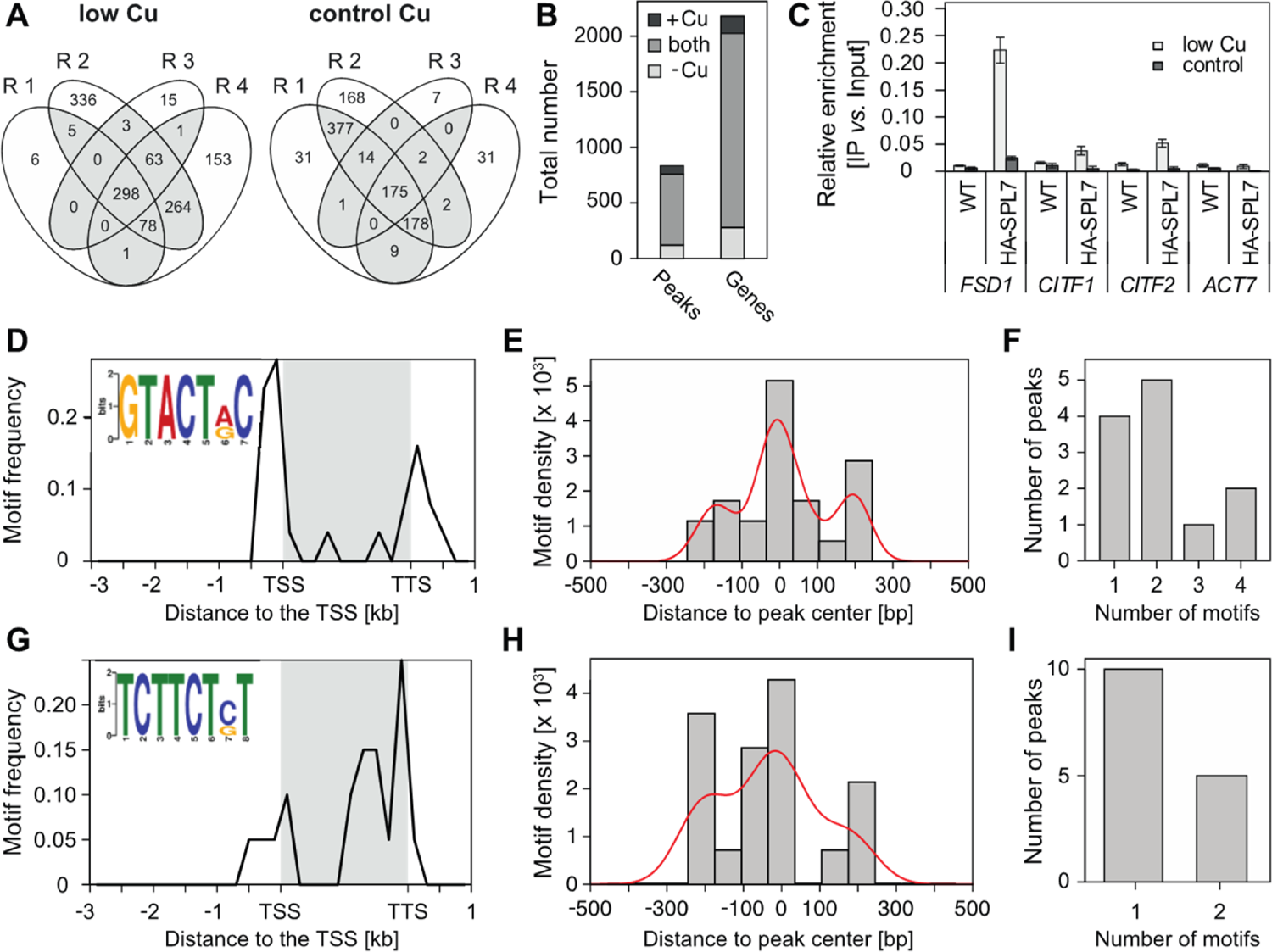
Genome-wide identification of SPL7 binding sites. (A) Venn diagrams showing the reproducibility of results. Given are the numbers of SPL7-binding segments (peaks) identified genome-wide by sequencing chromatin immunoprecipitates (ChIP-seq) in each of 4 independent experiments (as replicates, R). ChIP-seq was conducted on shoots of 21-d-old seedlings cultivated on control (0.5 µM Cu) and low-Cu (no Cu added) media supplemented with 1% (w/v) sucrose and solidified with EDTA-washed agar in vertically oriented glass petri dishes. (B) Bargraph showing the numbers of genomic SPL7-binding segments identified by ChIP-seq (shown on a grey background in A) and associated gene loci. (C) Validation of ChIP-seq results by ChIP-qPCR. Bargraph shows relative DNA enrichment of the promoter regions of novel candidates *FSD1*, *CITF1*, *CITF2* (*bHLH23*), and ACTIN7 (*ACT7*) as a negative control, quantified by ChIP-qPCR on immunoprecipitation (IP) and input samples. Chromatin was isolated from shoots of *spl7-1 P_SPL7_::HA-SPL7-HA:t_SPL7_* (HA-SPL7) and wild-type seedlings (WT, negative control) cultivated as described for (A). Input samples represent aliquots taken after chromatin shearing and before the addition of α-HA for the IP. See Supplemental Figure 12C, D for two additional independent experiments. (**D**-**I**) The GTACTRG motif (**D**-**F**, E-value = 8.20 x 10^-3^, 21 motif sites) and the TCTTCTST motif (**G**-**I**, E-value = 3.40 x 10^-2^, 19 motif sites) identified by MEME motif analysis. Motif frequency positional distribution (200 bp bin size), with conservation logo generated by MEME as an inset (**D, G**). Density plot summarizing distances of motif from center of peaks from ChIP-seq (75 bp bin size)(**E, H**). Bargraph showing motif copy numbers within single peaks (**F, I**). The gene body (grey box) of all corresponding genes was normalized to 2,000 bp (D, G). The red line visualizes the shape of the distribution (E, H). TSS: transcriptional start site, TTS: transcriptional termination site (positions from *A. thaliana* TAIR10 genome annotation).

In order to evaluate genomic SPL7 binding sites based on SPL7-dependence of transcript levels, we conducted RNA-seq on rosette tissues of wild-type and *spl7* seedlings cultivated as for ChIP-seq. Out of the top eight candidate genes for SPL7-dependent transcriptional activation identified through the largest *SPL7* dependence at the transcript level under low Cu conditions (Supplemental Data Set 3), five genes were also identified in our ChIP-seq under low Cu conditions: *FSD1*, bHLH transcription factor-encoding *bHLH160* (*COPPER-DEFICIENCY INDUCED TRANSCRIPTION FACTOR1*, *CITF1*; (Yan et al., 2017)), *bHLH23* (addressed here as *CITF2*), *YELLOW STRIPE-LIKE2* (*YSL2)* and *ZINC-REGULATED TRANSPORTER, IRON-REGULATED TRANSPORTER-RELATED PROTEIN2* (*ZIP2*) (Table 1, Supplemental Figure 12A-I, Supplemental Data Set 2). Of these, we tested three genes, *FSD1*, *CITF1* and *CITF2*, in independent ChIP-qPCR experiments and were able to confirm all three (Figure 7C, Supplemental Figure 12J and K). This supported a role of SPL7 in transcriptional activation under low-Cu conditions (Kropat et al., 2005).

**Table 1.**
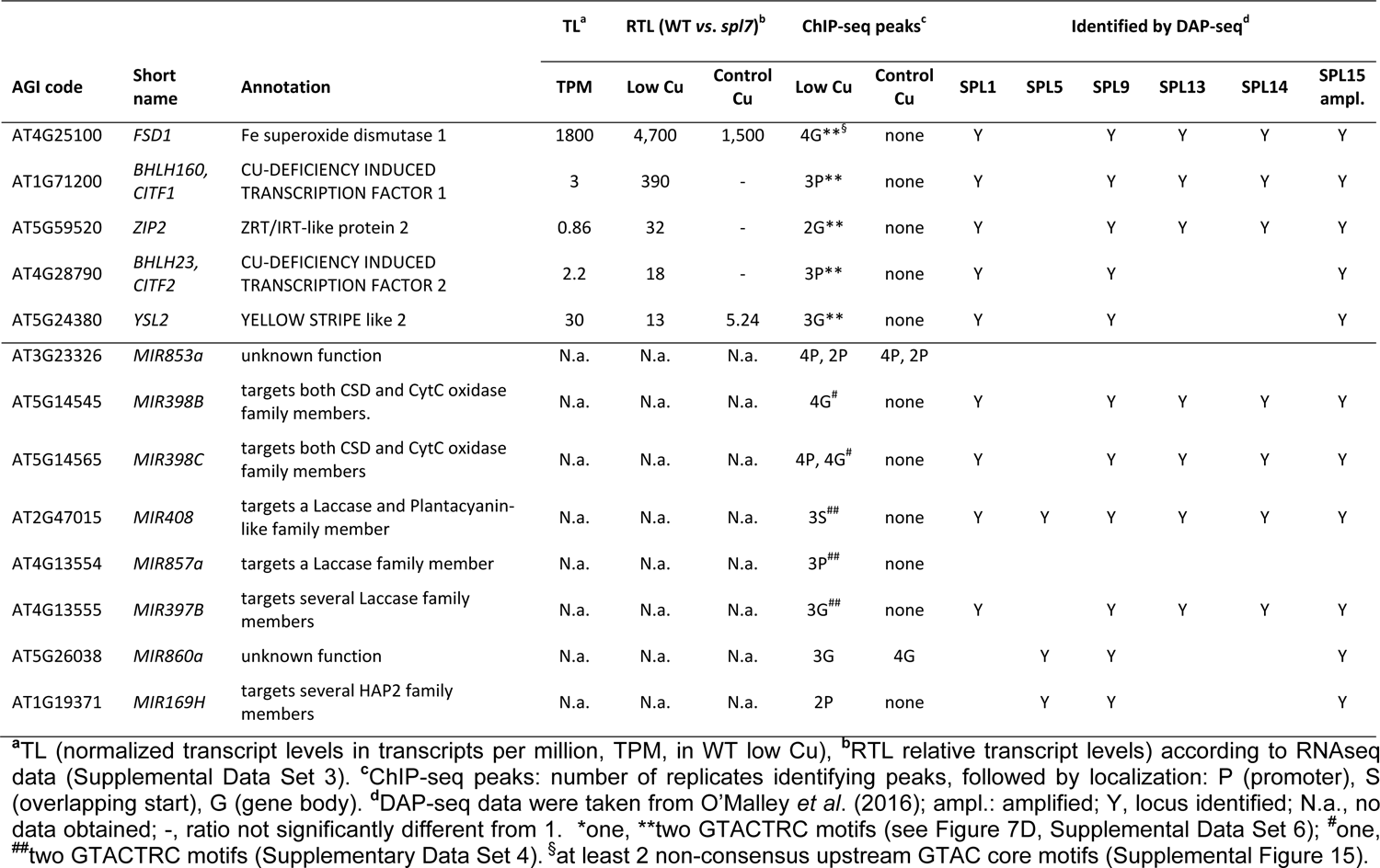
Genes undergoing maximal *SPL7*-dependent transcriptional regulation and microRNA loci associated with SPL7 binding sites according to ChIP-seq on shoot tissues of seedlings cultivated on copper-deficient medium.

All SPL family transcription factor proteins are thought to share the core GTAC motif of their DNA binding sites (Birkenbihl et al., 2005). Either individually or in common among genes previously identified by DAP-seq for SPL1, −5, −9, −13, −14 and −15 (O’Malley et al., 2016), the numbers of genes shared with our SPL7 ChIP-seq data were not significantly above expectations based on random picking (Supplemental Table 2, Supplemental Data Set 2). Notably, however, all of the top five genes highlighted above also contained binding sites for between three and five SPL proteins according to DAP-seq data (Table 1). This was also the case for the miRNA loci previously identified to be direct targets of SPL7 binding and SPL7-mediated transcriptional activation, miR408 and miR398B and C (Yamasaki et al., 2009; Zhang and Li, 2013).

Examining the sequences of all peaks identified by SPL7 ChIP-seq together, we could not identify any enriched sequence motif (Supplemental Data Set 4). Restricting our analysis to ChIP-seq peaks unique to low Cu, we identified the significantly enriched **GTAC**TA/GC motif that is partially identical to the previously reported A/T**GTAC**T/A, as well as significantly enriched AGGAAGC/T (reverse complement: A/G**CTTCCT**) that is reminiscent of TCT<u>TCTTCTC</u>**CTTCCT**C (Zhang et al., 2014)(Supplemental Figure 13). The subfamily II SPL protein IPA1 (SPL14) of *Oryza sativa* binds to differing core motifs, either GTAC or TGGGCC/T, dependent on environmental conditions (Wang et al., 2018). To further accommodate possible alternative SPL7 binding preferences dependent on Cu supply, we generated differing sets of peaks based on the transcriptional regulation of the associated genes for subsequent motif identification (Supplemental Data Sets 5 and 6). In this manner, we obtained 51 and 52 candidate genes for the direct activation and repression by SPL7 under low Cu conditions, respectively (Supplemental Data Set 5). Note that binding site positions predominantly upstream of transcriptional start sites were equally consistent with SPL7-dependent activation and repression in low-Cu conditions (compare Supplemental Figure 11D and F). Under control cultivation conditions, a clearly smaller number of 16 and 24 genes were candidates for undergoing direct SPL7-mediated activation and repression, respectively (Supplemental Data Set 5). SPL7 binding sites were positioned upstream or within the gene body at similar frequencies, suggesting either a lesser or a more complex role of SPL7 in transcriptional regulation under Cu-sufficient conditions (compare Supplemental Figure 11C and E).

We identified enriched motifs GTACTA/GC and AC/GAGAAGA (<u>TCTTCTC/G</u>T) among peaks identified by ChIP-seq associated with genes undergoing SPL7-dependent transcriptional activation, under low-Cu conditions (Figure 7D-I). Overall motif positioning, density and abundance supported the GTACRC motif more strongly than the TCTTCTST motif. Applying progressively relaxed filtering criteria for assembling sets of ChIP-seq peak-associated and *SPL7*-dependently regulated genes, identified multiple sequence motifs, part of which contain previously reported binding sites of other transcription factors (Supplemental Figure 14, Supplemental Data Sets 4 and 6). Four motifs newly identified here were predominantly positioned within gene bodies (Supplemental Figure 14A, D, G, I), similar to the other non-GTAC-containing motifs (see Figure 7G, Supplemental Figure 13D). Taken together, these results are consistent with a possible direct transcriptional activation by SPL7 under low Cu of *FSD1*, *CITF1* and *CITF2*, alongside a small set of other genes. Beyond this, our data are consistent with complex roles and interactions of SPL7 (Garcia-Molina et al., 2014a; Zhang et al., 2014; Yan et al., 2017).

Despite strong support for both positive and direct regulation of *FSD1* transcript levels by SPL7 (Table 1, Figures 6D and 7C, Supplemental Figures 4D and 12J and K), MEME did not identify *FSD1* to contain any of the enriched sequence motifs (Supplemental Data Sets 4 and 6). Manual inspection identified a total of 8 GTAC core motifs in the region corresponding to the *FSD1*-associated peak, with a minimum of a total of 4 copies of the GTAC core motif upstream of all annotated transcriptional start sites of *FSD1* (Supplemental Figure 15). Two of these copies correspond to GTACTRC, the motif identified here (see Figure 7D, Supplemental Figure 13A). There are two copies of AGTACA/TGTACT which was previously concluded to have the highest *in vitro* binding affinity for SPL7 out of several variants tested using electrophoretic mobility shift assays (Zhang et al., 2014).

## Discussion

### How do altered sugar responses arise in the *spl7* mutant?

In plant leaves, the largest cellular quota of both Fe (∼70% to 80%) and Cu (∼30%) are localized in chloroplasts (Shikanai et al., 2003; Yruela, 2013). The primary requirement for Cu in chloroplasts is in plastocyanin (Schubert et al., 2002). Cultivation of *Populus trichocarpa* in a Cu-limiting hydroponic solution caused a reduced abundance of PC protein and a strong decrease in photosynthetic electron transport rates, with plant biomass reduced by one half (Ravet et al., 2011). Arabidopsis plants lacking both plastocyanin isoforms are incapable of photoautotrophic growth and require exogenous sucrose to survive (Weigel et al., 2003). Biomass production was severely reduced in the *spl7-1* mutant cultivated under low-Cu conditions (Figure 1). It was plausible to attribute this growth defect to Cu limitation of photosynthesis in this mutant, given the known requirement for *SPL7* in the transcriptional activation of Cu acquisition and Cu economization processes (Yamasaki et al., 2009; Bernal et al., 2012). However, the unresponsiveness of growth of *spl7-1* to exogenous sucrose in low Cu and an enhanced accumulation of sugars in *spl7-1* implicated processes other than photosynthetic assimilate production in its growth defect (Figure 1, Supplemental Figures 2 to 4).

The second most relevant cellular site of Cu use in Arabidopsis is in the mitochondrial ETC, where Cu acts as a cofactor of cytochrome *c* oxidase (COX, complex IV). Total respiration was reduced by about 60% in Cu-deficient compared to Cu-replete cells of the green alga *Chlamydomonas reinhardtii* (Kropat et al., 2015). In wild-type *Arabidopsis thaliana*, electron flow *via* the COX-dependent cytochrome *c* pathway was not affected by Cu deficiency in the wild type (Figure 2), consistent with unaltered COX2 protein levels in *A. thaliana* and *P. trichocarpa* cultivated in low-Cu media (Yamasaki et al., 2007; Ravet et al., 2011). Interestingly, also in low-Cu grown *spl7-1*, there was no reduction in electron flux *via* the cytochrome *c* pathway. A slight *SPL7*-independent increase in transcript levels of *HCC1*, a chaperone involved in Cu delivery to COX (Attallah et al., 2011), under Cu deficiency may contribute to COX functionality in Cu-deficient Arabidopsis (Bernal et al., 2012). It is possible that *spl7-1* seedlings can only maintain adequate Cu supply to COX because of their attenuated growth (Bernal et al., 2012). An elevated total respiration rate in the *spl7-1* mutant under low Cu resulted from the activation of AOX-dependent respiration and may have a compensatory role in the presence of elevated ATP levels and NADH/NAD+ ratios in *spl7-1* (Figure 2, Supplemental Figure 6). An increased engagement of the alternative electron pathway in response to sugar stimuli is thought to constitute a balancing mechanism for an excess of reducing equivalents (Lambers, 1982; Azcón-Bieto et al., 1983). Dissimilar from the properties of COX-defective (Dahan et al., 2014) and other Cu-deficient genotypes (Figure 2D), we interpret enhanced respiration, increased NADH/NAD^+^ ratio, accumulation of ATP and an altered metabolite profile of *spl7-1* as consequences of an elevated sugar status, and not as its causes.

Given that we excluded major contributions from defects in the two most prominent Cu-dependent metabolic pathways, the attenuation of growth of *spl7-1* under low Cu despite high internal sugar levels might result from defective sugar/energy sensing or signaling. SnRK1 kinase activity is directly inhibited by T6P (Zhang et al., 2009; Zhai et al., 2018), a signaling metabolite of which we found elevated levels in *spl7-1* (Figures 2 and 5, Supplemental Figure 6). The responsiveness of downstream marker transcripts of the SnRK1 signaling pathway were consistent with its general functionality in *spl7-1* (Figure 3). The phosphorylation state of the TOR kinase target S6K was in accordance with sugar levels also in *spl7-1*, indicating that the TOR pathway is also generally functional in *spl7-1* under low Cu (Figures 1 and 4, Supplemental Figure 3 and 7). Different from these observations in *spl7-1*, TOR activity is decreased under nitrogen starvation (Liu et al., 2021). While TOR is involved in the phosphorylation of a variety of targets in several downstream signaling pathways, phosphorylation of S6K is responsive to sucrose and glucose (Xiong and Sheen, 2012; Dobrenel et al., 2016) and thus a suitable marker for TOR activity in our conditions. To summarize, TOR and SnRK1 activity responded to sugar signals irrespective of Cu availability and in both wild type and *spl7-1*, indicating against an involvement of these kinases in the *SPL7*-dependent integration of Cu deficiency responses with energy-dependent regulation of growth in Arabidopsis. Yet, we cannot exclude a possible partial attenuation of SnRK1 or TOR signaling in *spl7*.

Different from *spl7-1*, failure to grow as a consequence of impaired sulfur assimilation in the *A. thaliana sir1-1* mutant lacking the enzyme sulfite reductase was linked to reduced TOR activity (Dong et al., 2017). In agreement with sugar signals governing TOR activity, the *sir1-1* mutant contained reduced carbohydrate levels. Finally, transcript levels of published targets of nuclear Hexokinase1 (HXK1) signaling function *CAA* (*CA2*) and *CAB* (*LHCB1.1*) (Cho et al., 2006), as well as of the proposed downstream target of (HXK1)-dependent metabolic regulation, *NRT2.1* (Lejay et al., 2003; Lejay et al., 2008), were regulated as expected based on higher sugar contents in *spl7-1* (Supplemental Data Set 3; *CA2*, *LHCB1.1* four-fold down, *NRT1.2* eight-fold up in *spl7-1 vs.* WT under - Cu). The combination of Cu deficiency and the lack of *SPL7* might generate sink-limited conditions by affecting unknown processes downstream of the T6P/SnRK1 module. It was proposed that under sink-limited environmental conditions, namely low nitrogen supply or low temperature, the strongly inter-related levels of sucrose, T6P and SnRK1 activity are disconnected from the growth outcome (Nunes et al., 2013b). Regulator of G-protein Signaling 1 (RGS1) was proposed to operate as a plasma membrane-localized sensor of extracellular glucose (Urano et al., 2012). The activation of RGS1-dependent signaling was noted to require unexpectedly high extracellular glucose concentrations (Li and Sheen, 2016). A possible defect in RGS1-dependent signaling in *spl7* deserves attention in future work.

### *SPL7* and Cu deficiency in plant development

The levels of miR156, which post-transcriptionally represses subfamily II *SPL*s except *SPL8*, are progressively down-regulated with plant age in response to increasing sugar levels in leaves, thereby promoting vegetative phase change (Yang et al., 2011; Yang et al., 2013; Yu et al., 2013). The miR156/subfamily II SPL module was also implicated in the regulation of flowering time (Wu and Poethig, 2006; Schwarz et al., 2008; Wang et al., 2009). Transgenic plants containing reduced levels of the signaling sugar T6P contained reduced levels of *FLOWERING LOCUS T* transcript in whole rosettes, as well as elevated levels of miR156 in the shoot apex, and they flowered late (Wahl et al., 2013).

Our results are consistent with an attenuated response of developmental transitions to internal sugar levels in *spl7-1*. Compared to the wild type, flowering of *spl7-1* was delayed and unaltered by Cu supplementation, although at a given plant age the levels of the metabolizeable sugars sucrose and glucose as well as of T6P were increased (Figures 1 and 5, Supplemental Figures 3 and 8). Although remaining sugar-responsive in the *spl7-1* mutant, miR156 levels were strongly elevated in the mutant upon cultivation in low-sucrose-low-Cu conditions despite comparably high internal sugar levels, and miR156 could thus contribute to a condition-specific developmental delay (Figure 1 and Figure 5F). It is unlikely, however, that elevated levels of miR156 are causal in sugar accumulation of *spl7-1*, because sugar levels including T6P are not elevated in miR156-overexpressing plants (Ponnu et al., 2020). In *A. thaliana*, eight genetic loci encode miR156 precursors, and not all precursor transcripts decrease in abundance in response to sugars (Yang et al., 2013). Moreover, various abiotic stresses, such as heat stress, phosphate starvation, salt stress and drought, lead to increased miR156 levels (Hsieh et al., 2009; Cui et al., 2014; Stief et al., 2014).

Elevated Cu levels in aerial tissues compared to rosette leaves in the wild type indicated that Cu allocation to the inflorescence is prioritized, in agreement with the previously established role of Cu in plant fertility, for example in the Cu-binding protein plantacyanin involved in pollen tube guidance (Kim et al., 2003; Dong et al., 2005; Yan et al., 2017; Rahmati Ishka and Vatamaniuk, 2020). This prioritization was to a large extent dependent on *SPL7* (Figure 5B). Employing *in situ* RNA hybridization, we detected a signal for *SPL7* mRNA in the vasculature of the shoot apical meristem (Figure 5G). Future work will be required to analyze possible functions of SPL7 in the shoot apical meristem and during the reproductive phase of development.

### Genomic targets of SPL7 binding and transcriptional regulation

All SPL family proteins are characterized by a highly conserved recognition domain for the binding of a GTAC core DNA motif, and there is functional overlap among subfamily II SPLs in the regulation of plant development (Birkenbihl et al., 2005; Xing et al., 2010; Xing et al., 2013; Xu et al., 2016). Our phenotypic analysis of *spl7-1* mutants did not support a predominant functional antagonism between SPL7 and mir156-regulated subfamily II SPLs (Figure 5). Alternatively, direct or indirect target genes of regulation by the transcription factor SPL7 could include critical functions in sugar sensing, signaling or utilization. The combined analysis of genomic SPL7 binding sites and *SPL7*-dependent regulation of transcript levels did not reveal any evident well-characterized genes directly targeted by SPL7 and likely to cause the altered energy metabolism in *spl7-1* (Supplemental Data Set 5). Consequently, it is possible that SPL7 is required for the expression of a yet uncharacterized gene critical for sugar utilization.

The putative direct SPL7 target genes *CITF1* (*bHLH160*) and *CITF2* (*bHLH23*) identified here were reported as differentially regulated between the wild type and *spl7-2* in previous studies (Bernal et al., 2012; Yan et al., 2017). Additionally, our data support *FSD1* as a direct target of SPL7 (Table 1, Supplemental Figures 12 and 15). *FSD1* undergoes the most pronounced *SPL7*-dependent transcriptional regulation (Table 1, Supplemental Figure 4D). *FSD1* was also detected in the earlier ChIP-seq study of SPL7 (Zhang et al., 2014), together with the well-established direct targets of SPL7-dependent transcription under Cu deficiency, *MIR398B*, *MIR398C* and *MIR408* (Yamasaki et al., 2009; Zhang and Li, 2013). For all these genes, the positions of ChIP-seq peaks were consistent with SPL7 binding directly to the promoters under Cu deficiency (Table 1, Supplemental Figures 12 and 15). CITF1 was identified through its physical interaction with SPL7 and subsequently shown to be important in Cu deficiency responses of roots and reproductive organs, as well as for pollen fertility and jasmonate responses in flowers (Yan et al., 2017). It groups among bHLH transcription factors Ib, alongside the central Fe deficiency response regulatory transcription factors bHLH38/39/100 and 101 (Heim et al., 2003). Group VIIa of the bHLH transcription factors comprises CITF2 as well as the well-studied PHYTOCHROME INTERACTING FACOR and -LIKE bHLH proteins. Future work will address the functions of these transcription factors, as well as of superoxide dismutases, in particular of FSD1, in sugar responses and sugar utilization. An example of sugar signalling *via* reactive oxygen species was recently published (Roman et al., 2021).

Our ChIP-qPCR data suggested highly effective binding of SPL7 to *FSD1* in Cu-deficient conditions, associated with very high *FSD1* transcript levels, and – exceptionally – notable SPL7 binding even in Cu-sufficient seedlings, as well as both consensus and unusual GTAC-containing sequence elements in the promoter region (Figures 6 and 7, Table 1, Supplemental Figures 12, 13 and 15). Our observations support some residual SPL7 activity even when sufficient Cu is available, implying that small amounts of SPL7 are located inside the nucleus and active on a subset of target sites. Based on the transient expression of GFP-tagged SPL7 in leaves of *Nicotiana benthamiana*, it was proposed that SPL7 is anchored to the ER membrane by a transmembrane helix in its C-terminus and released into the cytoplasm only under Cu deficiency upon proteolytic cleavage at a site in the center of the protein. Accordingly, entry of the N-terminal half of SPL7, which includes the SBP domain, into the nucleus would then allow the transcriptional activation of Cu deficiency-responsive genes (Garcia-Molina et al., 2014b). Different from this proposed mechanism, immunoblots revealed only a single band corresponding approximately to the full protein size, irrespective of plant physiological Cu status, in a stably transformed *HA-SPL7-HA* line (Figure 6, Supplemental Figure 10).

Although we identified several motifs here, our data supported in a manner corresponding to simple expectations only the consensus GTACTRC motif in promoter regions for SPL7-mediated transcriptional enhancement primarily under low-Cu conditions (Table 1, Figure 7, Supplemental Figures 12 to 15). A shift in the SPL7 regulon under control Cu conditions, as exemplified by *FSD1*, could involve differing *cis*-regulatory DNA sequence elements that exhibit a higher binding affinity for SPL7, or alternatively as yet unidentified conditional protein interaction partners of SPL7. For example, KIN17 interacts with SPL7 specifically in aerial tissues and is involved in promoting Cu deficiency responses (Garcia-Molina et al., 2014a). SPL7 interacts with ELONGATED HYPOCOTYL (HY5) to enhance the levels of miR408 (Zhang et al., 2014). Physical interactions of SPL7 with other transcription factors may contribute to explaining the identification of multiple overrepresented motifs among SPL7 ChIP-seq peaks, the apparent binding of SPL7 within gene bodies under Cu-sufficient cultivation conditions, or the predominant localization of peaks in promoter regions also of genes repressed at the transcript level dependent on SPL7 (Figure 7G-I, Supplemental Figure 11C-F, 13D-F and 14, Supplemental Data Sets 4 and 6). Alternatively, SPL7 could act alone as a repressor of some of its direct target genes. To date, SPL7-dependent negative regulation was exclusively reported to occur indirectly *via* SPL7-dependent transcriptional activation of miRNA loci (Abdel-Ghany and Pilon, 2008; Yamasaki et al., 2009).

Another mechanism that can modulate the activity of SPL proteins was described for SPL14 in *Oryza sativa*, a homolog of AtSPL9/15 (Wang et al., 2018). The phosphorylation of a conserved serine residue in the SBP domain caused an altered DNA-binding specificity of OsSPL14, i.e. a change in preference for binding to a non-GTAC TGGGCC motif. OsSPL14 was thus found to have a dual role in alternatively promoting either yield or disease resistance. If a similar mechanism operated in SPL7, it could explain the additional identification of a non-GTAC SPL7-binding motif (Figure 7G-I, 13D-F and 14, Supplemental Data Set 6). Among genes associated with SPL7 ChIP-seq peaks, “response to hypoxia” was strongly overrepresented, but far less so among SPL7-dependently regulated transcripts (Supplemental Figure 16). CrCRR1 mediates transcriptional responses to both Cu deficiency and hypoxia (Hemschemeier et al., 2013). It now appears relevant to examine whether SPL7 and its orthologues regulate a subset of hypoxia responses in land plants, as well.

Future work will address the complex functions of SPL7 including the molecular mechanisms underlying growth and developmental impairment despite sugar accumulation in *spl7* mutants. Understanding the coordination of plant metal homeostasis with energy metabolism, growth and reproduction can help to increase crop yield and quality, especially on soils deficient in bioavailable Cu, which comprise more than 10% of the agricultural land in Europe (Reimann et al., 2014).

### Methods Plant material

*Arabidopsis thaliana* wild-type seeds (Col-0) were obtained from Lehle seeds (Round Rock, TX, USA). The *spl7-1* (SALK_093849) and *spl7-2* (SALK_125385) mutants are a T-DNA insertion lines from the Nottingham Arabidopsis Stock Centre and were characterized earlier (Yamasaki et al., 2009; Bernal et al., 2012). The generation of the *spl7-2 SPL7* (spl7-2_C) complemented line and transgenic *35S:amiRFRO4/FRO5* plants (line 27) was described in Bernal *et al*. (2012). The *paa1-3* loss-of-function mutant was a kind gift from Prof. Marinus Pilon (Shikanai et al., 2003), and *ran1-1* (N3808) was from the Nottingham Arabidopsis Stock Centre. Transgenic line *35S:amiR-TPS1* was kindly provided by Dr. Vanessa Wahl (Wahl et al., 2013). All mutants and transgenic lines are in the Col-0 genetic background. Primers used for genotyping are listed in Supplemental Table 3.

*HA-SPL7-HA* (*promSPL7::HA-SPL7-HA:termSPL7*) constructs were generated as follows (Lampropoulos et al., 2013). The *promSPL7* upstream region (−2506 to −5 from beginning of translational start codon) was PCR-amplified from genomic DNA *of A. thaliana* (Col-0) and cloned into the Greengate entry module pGGA000 *via* BsaI restriction. The genomic *SPL7* (AT5G18830.1) coding region (translational start to stop codon) was PCR-amplified and cloned into the pBluescript SK+ vector (Stratagene/Agilent Technologies, Waldbronn, Germany) which was used a PCR template for site-directed mutagenesis (A279T) to remove the internal BsaI recognition site in *SPL7* through a silent mutation. A Kozak consensus sequence and N-& C-terminal HA-tags were added to the genomic *SPL7* sequence with primer overhangs during the PCR amplification for the cloning into the Greengate entry module pGGI000 *via* BsaI restriction. The downstream *termSPL7* segment (+1 to +438 from end of translational stop codon) was PCR-amplified and cloned into the Greengate entry module pGGE000 *via* BsaI restriction. Using all entry modules and pGGF005 (Lampropoulos et al., 2013), the construct *promSPL7::HA-SPL7-HA:termSPL7:HygR* was assembled into the Greengate destination vector pGGZ003. The resulting binary plasmid was used to transform *Agrobacterium tumefaciens* (GV3130 [pSoup]), and the *spl7-1* mutant was transformed using the floral dip method (Clough and Bent, 1998). All primer sequences used for cloning are listed in Supplemental Table 3.

### Plant growth

Soil cultivation was in 16-h long days (145 µmol m^-2^ s^-1^, 22°C)/8-h night (18°C), with Cu conditions as described (Schulten et al., 2019). Glass petri dishes were soaked in 0.2 N HCl overnight and rinsed with deionized water to remove possible contaminant Cu before autoclaving. For plant cultivation in sterile culture on glass petri dishes, wild-type or mutant seeds were surface-sterilized, stratified in the dark at 4°C for 2 d and sown on a modified Hoagland solution (0.28 mM KH_2_PO_4_, 1.25 mM KNO_3_, 1.5 mM Ca(NO_3_)_2_, 0.75 mM MgSO_4_, 5 µM of a complex of Fe(III) and *N*,*N*′-di-(2-hydroxybenzoyl)-ethylenediamine-*N*,*N*′-diacetate (HBED), 25 µM H_3_BO_3_, 5 µM MnSO_4_, 5 µM ZnSO_4_, 0.5 µM CuSO_4_, 50 µM KCl, and 0.1 µM Na_2_MoO_4_, buffered at pH 5.7 with 3 mM 2-(*N*-morpholino)ethanesulfonate) in ultrapure water (Becher et al., 2004, with modifications), containing 1% (w/v) sucrose unless indicated otherwise and solidified with 1% (w/v) EDTA-washed Agar Type M (Sigma-Aldrich, Steinheim, Germany), as described (Schulten et al., 2019). Generally, 20 (or 40 for *spl7-1* under conditions without added CuSO_4_) seedlings were cultivated on each vertically orientated round glass petri dish (diameter of 150 mm) in 8-h short-day (145 µmol m^-2^ s^-1^, 22°C)/ 16-h night (18°C) cycles in a growth chamber (CFL Plant Climatics, Wertingen, Germany) for 21 d and pooled during harvest. For ChIP and RNA-seq, seedlings were grown for 21 d as described for Cu deficiency experiments in sterile culture, on glass petri dishes in 11-h short days (Schulten et al., 2019).

For experiments with combined Cu and sugar treatments, seedlings were pre-germinated on modified Hoagland’s medium without added CuSO_4_, containing 0.5% (w/v) sucrose and solidified with 1% (w/v) Agar Type M (Sigma-Aldrich, Steinheim, Germany), on square polypropylene petri dishes (120 mm x 120 mm) for 7 d. Seedlings were then transferred to controlled Cu growth conditions as described above for further cultivation for 14 d, with addition of either 0.05 µM CuSO_4_ (low Cu) or 0.5 µM CuSO_4_ (control conditions) and 0% or 6% (w/v) sucrose, respectively. All plants cultivated on soil and in agar-solidified media were harvested at Zeitgeber time (ZT) 3 h (3 h after lights on), or transferred darkness for 30 min at ZT 3 h for the quantification of ATP and NADH/NAD+, unless indicated otherwise.

For liquid cultures, 3.5 mg of surface-sterilized and stratified seeds of wild type and *spl7-1* were grown in 50 ml liquid 2x modified Hoagland solution with 0.5% (w/v) sucrose in 300 ml Erlenmeyer flasks placed on a rotation shaker (80 rpm) in an 8-h day (145 µmol m^-2^ s^-1^, 22°C)/16-h night (18°C) cycle in a growth chamber (CFL Plant Climatics, Wertingen, Germany) for 14 d. Note that all liquid cultures were germinated without added CuSO_4_ for one week, after which the medium was exchanged and half of the cultures of each genotype were cultivated with 1 µM CuSO_4_ for the remainder of the growth period. Two days before the sugar starvation treatment (d 13), the medium was exchanged again. The 3-h sugar starvation treatment was started at ZT 1 h (on d 15: Cultures were washed twice with sterile ultrapure water and fresh solutions as before without or with 0.5% (w/v) sucrose. For harvest (ZT 4 h), seedlings were washed in ultrapure water, shoots were separated from roots with a scalpel and blotted dry before snap-freezing in liquid nitrogen.

### Quantification of plant biomass and elemental concentrations

Quantification of plant biomass and elemental concentrations in plant tissues was conducted as described (Sinclair et al., 2017; Schulten et al., 2019). Aerial tissue samples were homogenized by grinding with a pestle in a mortar, which had been incubated in 0.2 N HCl overnight, rinsed in ultrapure water and dried at 60°C for > 1 h beforehand.

### RNA extraction and quantitative real time RT-PCR

RNA extraction, cDNA synthesis using oligo(dT)_18_ primers and reverse transcription quantitative real-time PCR (RT-qPCR) were performed as described (Schulten et al., 2019). Stem-loop pulsed reverse transcription of mature miRNAs was performed following a published protocol (Varkonyi-Gasic et al., 2007). Relative transcript levels (RTL) were calculated as follows: RTL = RE_m_^-ΔCT^, with RE_m_ as the mean of reaction efficiencies per primer pair and ΔC_T_ = C_T_(target gene) – C_T_(constitutively expressed reference genes: *EIF4*, *HEL* or *TUB3*), as described (Bernal et al., 2012). Primer sequences are listed in Supplemental Table 3.

### RNA in situ hybridization

Vegetative-stage shoot apical meristems were harvested from soil-grown plants at 8 d (cultivation in 16 h long days) and 30 d (cultivation in 8 h short days) of age, inflorescences from long-day grown plants at 15 to 20 cm height; tissues were processed to conduct RNA *in situ* hybridization, with probe synthesis from the cds of *SPL7* amplified and cloned into pGEMTeasy (Promega) according to manufacturer instructions, as described (Wahl et al., 2013).

### Metabolite extraction and measurement

Anthocyanins were extracted from aliquots (50 to 100 mg) of frozen ground shoot tissue powder in 2 ml methanol containing 1% (w/v) HCl by shaking overnight on a rotational shaker (150 rpm) at 4°C in the dark. Spectrophotometry was conducted on supernatants in 96-well plates and relative anthocyanin concentrations calculated as (A_530_ − 0.25*A_657_) g^-1^ fresh biomass (Rabino and Mancinelli, 1986).

Soluble sugars were extracted from aliquots (20 mg) of frozen ground shoot tissue powder in 250 µl of 80% (v/v) ethanol containing 10 mM HEPES/NaOH (pH 7) at 80°C for 30 min. After 10 min centrifugation at 3,500 rpm, the supernatant was collected and stored on ice. The extraction was repeated in 150 µl of the same solution and in 250 µl of 50% (v/v) ethanol containing 10 mM HEPES/NaOH (pH 7). Supernatants were combined and evaporated to dryness using a centrifugal vacuum dryer, followed by resuspension in 250 µl sterile ultrapure water. Aliquots (10 to 50 µl) were analyzed for sucrose and glucose contents with the K-SUFRG Kit (Megazyme, Bray, Ireland) in triplicate measurements in 96-well plates using a Synergy HTX microplate reader (BioTek, Bad Friedrichshall, Germany). For the quantification of starch, dried residues from the ethanolic extractions were washed in 1 ml ultrapure water, then resuspended in 400 µl of 0.1 M NaOH and heated at 98°C for 30 min. After pH neutralization, 100 µl starch degradation mix (16.8 units ml^-1^ amyloglucosidase [#10102857001; Roche, Mannheim, Germany] and 12 units ml^-1^ α-amylase [#10102814001; Roche, Mannheim, Germany] in 50 mM Na-acetate pH 4.9) was added per sample, followed by constant shaking at 37°C overnight. After centrifugation as described above, aliquots (50 µl) were used for the quantification of glucose levels as described above.

Trehalose 6-phosphate (T6P), phosphorylated intermediates and organic acids were extracted from aliquots (15 to 20 mg) of frozen ground shoot tissue powder in chloroform/methanol (3:7, v/v) and evaporated to dryness using a centrifugal vacuum dryer. The dried extracts were dissolved in 350 μl purified water and filtered through MultiScreen PCR-96 Filter Plate membranes (Merck Millipore, Darmstadt, Germany) to remove high-molecular-mass compounds. The metabolite extracts were subjected to high-performance anion-exchange chromatography coupled to tandem mass spectrometry (LC-MS/MS) as described (Lunn et al., 2006), with modifications (Figueroa et al., 2016).

For the quantification of adenosine triphosphate (ATP) levels, frozen ground shoot tissue (50 mg) was homogenized in 1 ml phenol (equilibrated with 10 mM TRIS/HCl, 1 mM EDTA, pH 8) and extracted into 0.5 ml of 10 mM TRIS/HCl (pH 8).a protocol for maize endosperm (Lappe et al., 2018). After centrifugation for 10 min at 10,000×*g,* the aqueous phase was extracted in an equal volume of chloroform. The supernatant was diluted 200-fold in 10 mM Tris/HCl (pH 8), and 100 μl sample volumes were assayed in triplicate for ATP content using the BacTiter-Glo bioluminescence kit (Promega, Mannheim, Germany). After 5 min incubation in the dark to allow for the decay of plate autofluorescence, photon emissions were recorded in white 96-well plates with an integration time of 0.5 s using a Synergy HTX microplate reader. ATP concentrations were quantified based on a standard curve (50 pM to 1 µM) and normalized to fresh biomass. For the quantification of NAD^+^ and NADH levels, 25-mg aliquots of frozen ground shoot tissues were extracted with 250 µl of either 0.2 M NaOH (NADH) or 0.2 M HCl (NAD^+^). The homogenates were centrifuged at 10,000xg and 4°C for 10 min. The supernatant (200 µl) was neutralized with 175 µl of either 0.2 M HCl or 0.2 M NaOH and the neutralization to pH 7 to 8 was confirmed with pH indicator paper. Aliquots of 50 µl were analyzed in triplicate for NAD^+^ and NADH with the NAD/NADH-Glo Kit (Promega, Mannheim, Germany) in white 96-well plates using a Synergy HTX microplate reader, with normalization to fresh biomass.

### Respiration measurements

The uptake of O_2_ by whole rosettes was quantified using a CG867 O_2_-meter (Schott Instruments, Weilheim, Germany) and the Clark-type oxygen electrode OX1100 (Schott Instruments, Weilheim, Germany), calibrated with saturated sodium dithionite solution and air-saturated water. Measurements were performed in darkness under constant stirring in a cuvette surrounded by a water-flow cooling system. Seedlings were kept in the dark for 30 min prior to the start of measurements to prevent photosynthesis. A total of 100 mg fresh biomass of rosette tissue was submerged in 3 mL of air-saturated 20 mM potassium phosphate buffer (pH 6.8) in the measuring cuvette. After 5 min of equilibration time, total respiration was monitored for 10 min during which O_2_ concentration was recorded every 30 sec. Then, potassium cyanide (KCN, 1 mM final concentration) or salicylhydroxamic acid (SHAM, 20 mM final concentration) were added to the cuvette for the measurement of cyanide (CN)- or SHAM-resistant respiration, respectively. Following the addition of an inhibitor and stabilization of the rate of O_2_ uptake during an equilibration time, respiration was monitored for another 7 to 10 min.

### Immunoblots

Immunological detection of S6K-1/2 and S6K-p and was performed as described with some modifications (Dong et al., 2017). In brief, total soluble protein was extracted from 50 mg of ground frozen shoot material in 250 µl 2× Laemmli buffer supplemented with 1% (v/v) phosphatase inhibitor cocktail 2 (Sigma-Aldrich, Steinheim, Germany). Proteins were denatured for 10 min at 90°C and separated using 10% (w/v) SDS-PAGE (22 mA, 2 h), followed by wet tank transfer to nitrocellulose membranes (100 V, 1 h, 4°C) (Towbin et al., 1979). After Ponceau staining (0.2% [w/v] Ponceau-S in 1% [v/v] acetic acid) to confirm equal loading per lane and blocking with 5% (w/v) BSA in TRIS-buffered saline containing 0.05% (v/v) Tween-20 (TBST) for 1 h, membranes were incubated with the primary antibody Phospho-p70 S6 Kinase (#9205; Cell Signaling, Frankfurt am Main, Germany) or anti-S6K1/2 (#AS12-1855; Agrisera, Vännas, Sweden) diluted 1:5,000 in TBST additionally containing 1% (w/v) BSA (TBSTB) at 4°C overnight. Membranes were washed 3 times 10 min in TBST and then incubated with an HRP-conjugated secondary antibody diluted 1:15,000 in TBSTB at RT for 1 h. After repeating the washes with TBST, detection was carried out with Pierce ECL Western Blotting Substrate (ThermoFisher, Schwerte, Germany) using a Fusion Fx7 GelDoc (Vilber Lourmat, Eberhardzell, Germany).

For the immunological detection of HA epitope-tagged SPL7, total soluble proteins were extracted from 50 mg of ground frozen shoot material with 100 µl 2× Laemmli buffer (#S3401, Sigma-Aldrich, Steinheim, Germany). Proteins were denatured and separated as described above, with wet/tank transfer overnight (60 mA, 4°C) (Towbin et al., 1979). After Ponceau staining of the membrane as described above and blocking with 5% (w/v) blotting-grade milk powder in TBST for 1 h, membranes were incubated with the primary anti-HA antibody (#26183; ThermoFisher, Schwerte, Germany) diluted 1:5,000 in TBST containing 1% (w/v) milk powder (TBSTM) or anti-Actin antibody diluted 1:2,500 in TBSTM (#AS132640; Agrisera, Vännas, Sweden) at RT for 2 h. Membranes were washed as described above and then incubated with HRP-conjugated secondary antibodies (ThermoFisher, Schwerte, Germany) diluted 1:250 (#32430 following anti-HA) in TBSTM or 1:2,500 (#31466, following anti-Actin) in TBSTM at RT for 1 h, followed by washing as described above and detection with ECL Select Western Blotting Reagent (GE Healthcare, Little Chalfont, England) as described above.

### Flowering time parameters

The number of days to flowering was counted from the day the seeds were placed in the growth chamber after stratification until the bolting shoot reached ∼0.5 cm in length. Rosette leaves were counted alongside plant growth until the bolting shoot had reached ∼0.5 cm in length. Plants were photographed 10 d after bolting to record the flowering phenotype. Rosette leaves and aerial tissues above the rosette were then harvested for analysis of elemental contents.

### Chromatin immunoprecipitation and analysis

For samples of pooled rosette leaves of 80 *spl7-1 HA-SPL7-HA* or wild-type seedlings grown on petri plates, chromatin was crosslinked by vacuum-infiltrating seedlings in a solution of 1% (w/v) formaldehyde in PBS buffer for 5 min, briefly releasing the vacuum, followed by vacuum-infiltration for another 10 min (Gendrel et al., 2002). The cross-linking reaction was stopped by adding glycine to a final concentration of 0.125 M and vacuum-infiltrating for 5 min. The rosettes were rinsed twice with sterile ultrapure water (Purelab Flex 2; ELGA LabWater, Celle, Germany), blotted dry and snap-frozen in liquid nitrogen. After grinding the plant material to a fine powder with a mortar and pestle in liquid nitrogen, nuclei were isolated as described (Moehs et al., 1988), with all extraction buffers additionally containing 1 mM phenylmethylsulfonyl fluoride (PMSF) and protease inhibitor cocktail (PIC) in a 1:1000 dilution (P9599, Sigma-Aldrich, Steinheim, Germany). Chromatin was extracted with nuclei lysis buffer (50 mM TRIS/HCl pH 8, 10 mM EDTA, 1% [w/v] SDS, 1 mM PMSF, PIC) and sonicated using the Bioruptor Pico (Diagenode, Seraing, Belgium) for 5 cycles (30 s on/ 30 s off) to achieve an average fragment size of 200 bp. After removing cellular debris by centrifugation (twice for 10 min at 10.000 x *g*), the chromatin was diluted 10-fold with ChIP dilution buffer (1.1% [v/v] Triton x-100, 1.2 mM EDTA, 16.7 mM TRIS/HCl pH 8, 167 mM NaCl) and pre-cleared at 4°C for 1 h by incubating with 80 µl of protein A beads (#17127901, GE Healthcare, Freiburg, Germany), pre-equilibrated with ChIP dilution buffer. After the pre-clear, an aliquot corresponding to 1% (v/v) of the starting chromatin volume was removed for use as the input DNA control. Monoclonal anti-HA antibody 12CA5 (ROAHA; Roche, Mannheim, Germany) was used to immunoprecipitate HA-SPL7-HA-bound chromatin (5 µg of antibody per IP) with 50 µl of pre-equilibrated protein A beads at 4°C overnight on a rotator (12 rpm). Washes of the immunocomplexes were performed as follows: one wash step with low salt wash buffer (150 mM, 0.1% (w/v) SDS, 1% (v/v) Triton X-100, 2 mM EDTA, 20 mM TRIS/HCl pH 8), two wash steps with high salt wash buffer (500 mM, 0.1% (w/v) SDS, 1% (v/v) Triton X-100, 2 mM EDTA, 20 mM TRIS/HCl pH 8), three wash steps with LiCl wash buffer (250 mM LiCl, 0.5% (w/v) IGEPAL, 0.5% (w/v) sodium deoxycholate, 1 mM EDTA, 10 mM TRIS/HCl pH 8) and two wash steps with TE buffer (1 mM EDTA, 10 mM TRIS/HCl pH 8) (Yamaguchi et al., 2014a, with modifications). The immunocomplexes were eluted twice from the protein A beads with freshly prepared elution buffer (1% [w/v] SDS, 0.1 M NaHCO_3_) by incubating at 65°C for 15 min at 1200 rpm in a ThermoMixer Comfort (Eppendorf AG, Hamburg, Germany). NaCl was added to the eluates and the input DNA aliquots to a final concentration of 0.2 M; samples were then incubated overnight (65°C, 600 rpm) for de-crosslinking and treated with proteinase K at 42°C for 1 h. DNA was purified with the NucleoSpin PCR Cleanup kit (#740609; Macherey-Nagel, Düren, Germany) with buffer NTB (#740595.150; Macherey-Nagel, Düren, Germany) and eluted in 15 µl of 5 mM TRIS/HCl pH 8.5.

ChIP samples were tested for enrichment by qPCR, performed as described, measuring enrichment on the promoter regions of *FSD1* and *MIR408* as positive controls and *ACTIN7* as a negative control (Schulten et al., 2019). The sequences of the primers used in ChIP-qPCR are listed in Supplemental Table 1. Libraries for ChIP-seq were prepared and sequenced at the Max-Planck-Genome Center Cologne on an Illumina HiSeq3000 instrument (Romera-Branchat et al., 2020).

### Transcriptome sequencing

Wild-type and *spl7-1* seedlings were grown and harvested as described above for ChIP. Total RNA was extracted from 50-mg aliquots of frozen ground rosette tissues using the RNeasy Plant Mini Kit (#74904, Qiagen, Hilden, Germany) including on-column DNase digestion to remove any contaminating genomic DNA (#79254, Qiagen, Hilden, Germany). RNA was quantified using the Qubit RNA Assay Kit in a Qubit 2.0 Fluorometer (Thermo Fisher, Schwerte, Germany). RNA integrity was assessed with the RNA Nano 6000 Assay Kit of the Agilent Bioanalyzer 2100 system (Agilent Technologies, Waldbronn, Germany) and the RNA integrity number (RIN) was between 8 and 9 for all samples. Library construction and next-generation sequencing were performed using Illumina technology by the Novogene Company (Wan Chai, Hong Kong). In brief, 1 µg of total RNA was used as input for library preparation, mRNA was enriched with oligo(dT)-beads and cDNA was synthesized with random hexamer primers with the NEB Next Ultra RNA Library Prep Kit (NEB, Ipswich, USA). All libraries were sequenced using the Illumina NovaSeq6000 platform in paired-end mode with a read length of 150 bp.

### Chromatin immunoprecipitation sequence data analysis

Upon sequencing of chromatin immonoprecipitates and the corresponding input control libraries of all samples from four independent experiments (addressed as four replicates below), adapter sequences were removed and low-quality ends were trimmed from raw reads using cutadapt (Martin, 2011) and Trimmomatic (Bolger et al., 2014), respectively. Read length distribution was summarized using the density function available in R (R Core Team, 2019). Reads were mapped to the *A. thaliana* TAIR10 reference genome assembly using bowtie2 (Langmead et al., 2009; Langmead and Salzberg, 2012) with default settings and a minimum mapping quality of 30, obtaining between 9 and 11 mio. reads per sample (median 10.4 mio. reads). ChIP peaks were identified using MACS2 v2.1.3.3 (Zhang et al., 2008) on each IP together with the respective INPUT from each sample by adjusted manifold confidence of enrichment ratio (-m 2 20; FDR ≤ 0.05 (Benjamini and Hochberg, 1995). The peaks of replicates were merged when at least 80% of the length of the shortest genomic segment covered by a peak overlapped with the segment covered by another peak (Romera-Branchat et al., 2020). A merged peak segment was thus at least as long as, or longer than, the broadest of the replicate peaks merged. A peak was considered as present in both low Cu and control Cu conditions when there was any overlap between the segments defined by peak center (mid-point of segment covered by the merged peak) ± 30% length of segment covered by the merged peak, respectively. Peaks were associated with *A. thaliana* genes (TAIR10) using Bioconductor R package chipPeakAnno and as previously described (Romera-Branchat et al., 2020).

### RNAseq data analysis

Adapters and low-quality bases were removed in RNAseq data from three independent experiments using trimmomatic (Bolger et al., 2014), keeping reads of at least 120 bp in length (19.6 to 27.3 million trimmed reads per sample). Reads were mapped to the *A. thaliana* TAIR10 genome assembly using hisat2 v2.1, excluding unaligned reads from the output BAM file (Lamesch et al., 2012; Berardini et al., 2015; Kim et al., 2019), followed by corrections of multiple mapping with identical start or end positions through COMEX 2.1 (Pietzenuk et al., 2016). Total numbers of counts per gene were retrieved non-strand specifically using Qualimap v.2.2.1 employing the proportional algorithm for multiply mapping reads and the *A. thaliana* Araport11 genome annotation (Okonechnikov et al., 2016; Cheng et al., 2017). Differentially expressed genes were identified using DESeq2 version 3.11 on the R 4.0.3 statistical computing platform with default settings using the local fit type (fitType = “local”) (Love et al., 2014; R Core Team, 2019)(Supplemental Data Sets 3 and 5).

### Motif discovery

For each set of ChIP peaks (Supplemental Data Set 1) passing a given set of filters (Supplemental Data Sets 3 to 6), the corresponding genomic sequences were extracted and analyzed for enriched motifs using MEME-ChIP (Bailey et al., 2009; Machanick and Bailey, 2011), with the following parameters: maximum motif width (-meme-maxw) 10, minimum motif width (-meme-minw) 4, motif occurrences mode (-meme-mod), anr (any number of repetitions), and motif database (-db) ArabidopsisDAPv1.meme. The predicted motifs were extracted from the corresponding FIMO output GFF file using a custom-made shell script and subsequently collated with the corresponding set of genes for calculation of motif distance to peak center, motif distance to TSS (gene transcriptional start site) and motif incidence using custom-made scripts.

### Quantification and statistical analyses

Each experiment was repeated independently two to three times. Poorly germinated seedlings or obvious phenotypic outlier individuals were excluded from harvest and measurements. Data are shown from one representative experiment, with *n* as indicated in the figure legends. Statistical analyses were performed with R v.3.5.0 (R Core Team, 2019). ANOVA followed by a Tukey’s post hoc test (*p* < 0.05) was conducted for datasets with homoscedasticity and a normal distribution of residuals. For all other datasets, pairwise *t*-tests (Student’s *t*-test or Welch *t*-test as applicable) with false discovery rate adjustment (*q*-value < 0.05) were performed instead (Storey et al., 2019).

## Accession numbers

Sequence data from this article can be found at ENA, EMBL-EBI under accession number PRJEB47134 (ERR6548266 - ERR6548277 for RNA-seq, ERR6558497 - ERR6558513 for ChIP-seq).

## Acknowledgements

We thank Petra Düchting (Ruhr University Bochum, Germany) for multi-element analysis, John E. Lunn (Max Planck Institute of Molecular Plant Physiology, Golm, Germany) for discussions of metabolite data as well as Dr. Yihan Dong (Institute of Molecular Biology of Plants, Strasbourg, France) and Dr. Markus Wirtz (Centre for Organismal Studies, Heidelberg, Germany) for advice on S6K immunoblots. This work was funded by the Deutsche Forschungsgemeinschaft (Kr1967/15-1) and Ruhr University Bochum, Germany.

## Author contributions

A.S., J.Q., M. R.-B., R.F. and V.W. performed experiments, A.S., B.P., M.K., E.S., G.C. and U.K. conducted computational or other data analysis, A.S. and U.K. designed the research and wrote the manuscript, with contributions from J.Q. and B.P., all authors edited the manuscript.

## Supplemental Data

Supplemental Table 1. Metabolite data related to Fig. 3D and Fig. S6C-D

Supplemental Table 2. Number of genes commonly identified in the present study and in earlier studies.

Supplemental Table 3. Oligonucleotides used in this study

Supplemental Figure 1. Effects of Cu deficiency and sucrose on nutrient metal concentrations in wild-type and *spl7-1* mutant seedlings.

Supplemental Figure 2. Sucrose does not stimulate shoot fresh biomass gain in the *spl7-1* mutant cultivated under low-Cu conditions.

Supplemental Figure 3. Independent experiment (repeat) related to Figure 1 (C-E).

Supplemental Figure 4. Effects of Cu deficiency and sucrose on relative transcript levels of known *SPL7*-dependently expressed genes.

Supplemental Figure 5. Starch levels in wild-type and *spl7-1* mutant seedlings upon cultivation in agar-solidified media containing different combinations of Cu and sucrose.

Supplemental Figure 6. Independent experiments (repeats) related to Figure 2.

Supplemental Figure 7. Two independent experiments (repeats) related to Figure 4.

Supplemental Figure 8. Two independent experiments (repeats) related to Figure 5 (D, E).

Supplemental Figure 9. Complementation of the *spl7-1* mutant through the *P_SPL7_::HA-SPL7-HA:t_SPL7_* transgene (two independent lines in addition to line 4-1 shown in Figure 6).

Supplemental Figure 10. Full image and independent replication of immunoblot shown in Figure 6C.

Supplemental Figure 11. Distribution of genomic SPL7 binding sites relative to genes.

Supplemental Figure 12. SPL7 binding profiles at chosen loci, and two independent repeats related to Figure 7C.

Supplemental Figure 13. Enriched motifs identified by MEME motif analysis among SPL7 binding sites detected exclusively under low Cu.

Supplemental Figure 14. Enriched motifs identified by MEME motif analysis on subgroups of SPL7 binding sites delineated by including information on gene expression.

Supplemental Figure 15. Putative SPL7-binding motifs at the *FE SUPEROXIDE DISMUTASE1* (*FSD1*) locus identified by ChIP-seq.

Supplemental Figure 16. GO and KEGG enrichment analyses.

**Supplemental Data Set 1.** Genomic binding sites of SPL7 detected by ChIP-seq.

**Supplemental Data Set 2.** Genes associated with SPL7 binding sites detected by ChIP-seq, and comparison with published DAP-seq data for other Arabidopsis SPL proteins.

**Supplemental Data Set 3.** Universal RNA-seq data.

**Supplemental Data Set 4.** Motifs in genomic SPL7 binding sites identified by ChIP-seq.

**Supplemental Data Set 5.** Genes exhibiting both *SPL7*-dependent regulation of transcript abundance and SPL7-binding peaks identified by ChIP-seq.

**Supplemental Data Set 6.** Putative SPL7-binding sequence motifs within the SPL7-binding region at the *FE SUPEROXIDE DISMUTASE1* (*FSD1*) locus identified by ChIP-seq.

## References

1. Abdel-Ghany, S.E., and Pilon, M. (2008). MicroRNA-mediated systemic down-regulation of copper protein expression in response to low copper availability in Arabidopsis. J Biol Chem 283, 15932–15945.

2. Abdel-Ghany, S.E., Müller-Moulé, P., Niyogi, K.K., Pilon, M., and Shikanai, T. (2005). Two P-type ATPases are required for copper delivery in *Arabidopsis thaliana* chloroplasts. Plant Cell 17, 1233–1251.

3. Assunção, A.G.L., Herrero, E., Lin, Y.-F., Huettel, B., Talukdar, S., Smaczniak, C., Immink, R.G.H., van Eldik, M., Fiers, M., Schat, H., and Aarts, M.G.M. (2010). *Arabidopsis thaliana* transcription factors bZIP19 and bZIP23 regulate the adaptation to zinc deficiency. Proc Natl Acad Sci USA 107, 10296.

4. Attallah, C.V., Welchen, E., Martin, A.P., Spinelli, S.V., Bonnard, G., Palatnik, J.F., and Gonzalez, D.H. (2011). Plants contain two SCO proteins that are differentially involved in cytochrome *c* oxidase function and copper and redox homeostasis. J Exp Bot 62, 4281–4294.

5. Azcón-Bieto, J., Lambers, H., and Day, D.A. (1983). Effect of photosynthesis and carbohydrate status on respiratory rates and the involvement of the alternative pathway in leaf respiration. Plant Physiol 72, 598–603.

6. Baena-Gonzalez, E., and Hanson, J. (2017). Shaping plant development through the SnRK1-TOR metabolic regulators. Curr Opin Plant Biol 35, 152–157.

7. Baena-González, E., Rolland, F., Thevelein, J.M., and Sheen, J. (2007). A central integrator of transcription networks in plant stress and energy signalling. Nature 448, 938.

8. Bahr, J.T., and Bonner, W.D., Jr. (1973). Cyanide-insensitive respiration. II. Control of the alternate pathway. J Biol Chem 248, 3446–3450.

9. Bailey, T.L., Boden, M., Buske, F.A., Frith, M., Grant, C.E., Clementi, L., Ren, J., Li, W.W., and Noble, W.S. (2009). MEME SUITE: tools for motif discovery and searching. Nucleic Acids Res 37, W202–208.

10. Becher, M., Talke, I.N., Krall, L., and Krämer, U. (2004). Cross-species microarray transcript profiling reveals high constitutive expression of metal homeostasis genes in shoots of the zinc hyperaccumulator *Arabidopsis halleri*. Plant J 37, 251–268.

11. Benjamini, Y., and Hochberg, Y. (1995). Controlling the False Discovery Rate: A Practical and Powerful Approach to Multiple Testing. Journal of the Royal Statistical Society: Series B (Methodological) 57, 289–300.

12. Berardini, T.Z., Reiser, L., Li, D., Mezheritsky, Y., Muller, R., Strait, E., and Huala, E. (2015). The Arabidopsis information resource: Making and mining the “gold standard” annotated reference plant genome. Genesis 53, 474–485.

13. Bernal, M., Casero, D., Singh, V., Wilson, G.T., Grande, A., Yang, H., Dodani, S.C., Pellegrini, M., Huijser, P., Connolly, E.L., Merchant, S.S., and Krämer, U. (2012). Transcriptome sequencing identifies *SPL7*-regulated copper acquisition genes *FRO4/FRO5* and the copper dependence of iron homeostasis in Arabidopsis. Plant Cell 24, 738–761.

14. Birkenbihl, R.P., Jach, G., Saedler, H., and Huijser, P. (2005). Functional dissection of the plant-specific SBP-domain: overlap of the DNA-binding and nuclear localization domains. J Mol Biol 352, 585–596.

15. Blaby-Haas, C.E., Padilla-Benavides, T., Stube, R., Argüello, J.M., and Merchant, S.S. (2014). Evolution of a plant-specific copper chaperone family for chloroplast copper homeostasis. Proc Natl Acad Sci USA 111, E5480–5487.

16. Bolger, A.M., Lohse, M., and Usadel, B. (2014). Trimmomatic: a flexible trimmer for Illumina sequence data. Bioinformatics 30, 2114–2120.

17. Cardon, G., Hohmann, S., Klein, J., Nettesheim, K., Saedler, H., and Huijser, P. (1999). Molecular characterisation of the Arabidopsis SBP-box genes. Gene 237, 91–104.

18. Caspar, T., Huber, S.C., and Somerville, C. (1985). Alterations in Growth, Photosynthesis, and Respiration in a Starchless Mutant of *Arabidopsis thaliana* (L.) Deficient in Chloroplast Phosphoglucomutase Activity. Plant Physiol 79, 11–17.

19. Chao, L.M., Liu, Y.Q., Chen, D.Y., Xue, X.Y., Mao, Y.B., and Chen, X.Y. (2017). Arabidopsis Transcription Factors SPL1 and SPL12 Confer Plant Thermotolerance at Reproductive Stage. Mol Plant 10, 735–748.

20. Cheng, C.Y., Krishnakumar, V., Chan, A.P., Thibaud-Nissen, F., Schobel, S., and Town, C.D. (2017). Araport11: a complete reannotation of the *Arabidopsis thaliana* reference genome. Plant J 89, 789–804.

21. Cho, Y.H., Yoo, S.D., and Sheen, J. (2006). Regulatory functions of nuclear hexokinase1 complex in glucose signaling. Cell 127, 579–589.

22. Clemens, S. (2001). Molecular mechanisms of plant metal tolerance and homeostasis. Planta 212, 475–486.

23. Clough, S.J., and Bent, A.F. (1998). Floral dip: a simplified method for *Agrobacterium*-mediated transformation of *Arabidopsis thaliana*. Plant J 16, 735–743.

24. Colangelo, E.P., and Guerinot, M.L. (2004). The Essential Basic Helix-Loop-Helix Protein FIT1 Is Required for the Iron Deficiency Response. Plant Cell 16, 3400.

25. Cui, L.-G., Shan, J.-X., Shi, M., Gao, J.-P., and Lin, H.-X. (2014). The miR156-SPL9-DFR pathway coordinates the relationship between development and abiotic stress tolerance in plants. Plant J 80, 1108–1117.

26. Dahan, J., Tcherkez, G., Macherel, D., Benamar, A., Belcram, K., Quadrado, M., Arnal, N., and Mireau, H. (2014). Disruption of the *CYTOCHROME C OXIDASE DEFICIENT1* gene leads to cytochrome *c* oxidase depletion and reorchestrated respiratory metabolism in Arabidopsis. Plant Physiol 166, 1788–1802.

27. Deprost, D., Yao, L., Sormani, R., Moreau, M., Leterreux, G., Nicolaï, M., Bedu, M., Robaglia, C., and Meyer, C. (2007). The Arabidopsis TOR kinase links plant growth, yield, stress resistance and mRNA translation. EMBO Rep 8, 864–870.

28. Dobrenel, T., Mancera-Martínez, E., Forzani, C., Azzopardi, M., Davanture, M., Moreau, M., Schepetilnikov, M., Chicher, J., Langella, O., Zivy, M., Robaglia, C., Ryabova, L.A., Hanson, J., and Meyer, C. (2016). The Arabidopsis TOR Kinase Specifically Regulates the Expression of Nuclear Genes Coding for Plastidic Ribosomal Proteins and the Phosphorylation of the Cytosolic Ribosomal Protein S6. Front Plant Sci 7, 1611–1611.

29. Dong, J., Kim, S.T., and Lord, E.M. (2005). Plantacyanin Plays a Role in Reproduction in Arabidopsis. Plant Physiol 138, 778.

30. Dong, Y., Silbermann, M., Speiser, A., Forieri, I., Linster, E., Poschet, G., Allboje Samami, A., Wanatabe, M., Sticht, C., Teleman, A.A., Deragon, J.M., Saito, K., Hell, R., and Wirtz, M. (2017). Sulfur availability regulates plant growth via glucose-TOR signaling. Nat Commun 8, 1174.

31. Figueroa, C.M., and Lunn, J.E. (2016). A Tale of Two Sugars: Trehalose 6-Phosphate and Sucrose. Plant Physiol 172, 7–27.

32. Figueroa, C.M., Feil, R., Ishihara, H., Watanabe, M., Kölling, K., Krause, U., Höhne, M., Encke, B., Plaxton, W.C., Zeeman, S.C., Li, Z., Schulze, W.X., Hoefgen, R., Stitt, M., and Lunn, J.E. (2016). Trehalose 6–phosphate coordinates organic and amino acid metabolism with carbon availability. Plant J 85, 410–423.

33. Foster, A.W., Osman, D., and Robinson, N.J. (2014). Metal Preferences and Metallation. J Biol Chem 289, 28095–28103.

34. Fraústo da Silva, J.J.R., and Williams, R.J.P. (2001). The Biological Chemistry of the Elements. (Oxford, UK: Oxford University Press).

35. Garcia-Molina, A., Xing, S., and Huijser, P. (2014a). A conserved KIN17 curved DNA-binding domain protein assembles with SQUAMOSA PROMOTER-BINDING PROTEIN-LIKE7 to adapt Arabidopsis growth and development to limiting copper availability. Plant Physiol 164, 828–840.

36. Garcia-Molina, A., Xing, S., and Huijser, P. (2014b). Functional characterisation of Arabidopsis SPL7 conserved protein domains suggests novel regulatory mechanisms in the Cu deficiency response. BMC Plant Biol 14, 231.

37. Garcia, L., Welchen, E., Gey, U., Arce, A.L., Steinebrunner, I., and Gonzalez, D.H. (2016). The cytochrome c oxidase biogenesis factor AtCOX17 modulates stress responses in Arabidopsis. Plant Cell Environ 39, 628–644.

38. Gendrel, A.-V., Lippman, Z., Yordan, C., Colot, V., and Martienssen, R.A. (2002). Dependence of Heterochromatic Histone H3 Methylation Patterns on the Arabidopsis Gene *DDM1*. Science 297, 1871–1873.

39. Gibon, Y., Bläsing, O.E., Palacios-Rojas, N., Pankovic, D., Hendriks, J.H., Fisahn, J., Hohne, M., Gunther, M., and Stitt, M. (2004). Adjustment of diurnal starch turnover to short days: depletion of sugar during the night leads to a temporary inhibition of carbohydrate utilization, accumulation of sugars and post-translational activation of ADP-glucose pyrophosphorylase in the following light period. Plant J 39, 847–862.

40. He, J., Xu, M., Willmann, M.R., McCormick, K., Hu, T., Yang, L., Starker, C.G., Voytas, D.F., Meyers, B.C., and Poethig, R.S. (2018). Threshold-dependent repression of *SPL* gene expression by miR156/miR157 controls vegetative phase change in *Arabidopsis thaliana*. PLOS Genet 14, e1007337.

41. Heim, M.A., Jakoby, M., Werber, M., Martin, C., Weisshaar, B., and Bailey, P.C. (2003). The basic helix-loop-helix transcription factor family in plants: a genome-wide study of protein structure and functional diversity. Mol Biol Evol 20, 735–747.

42. Heineke, D., Riens, B., Grosse, H., Hoferichter, P., Peter, U., Flügge, U.I., and Heldt, H.W. (1991). Redox Transfer across the Inner Chloroplast Envelope Membrane. Plant Physiol 95, 1131–1137.

43. Hemschemeier, A., Casero, D., Liu, B., Benning, C., Pellegrini, M., Happe, T., and Merchant, S.S. (2013). Copper response regulator1-dependent and -independent responses of the Chlamydomonas reinhardtii transcriptome to dark anoxia. Plant Cell 25, 3186–3211.

44. Hsieh, L.-C., Lin, S.-I., Shih, A.C.-C., Chen, J.-W., Lin, W.-Y., Tseng, C.-Y., Li, W.-H., and Chiou, T.-J. (2009). Uncovering Small RNA-Mediated Responses to Phosphate Deficiency in Arabidopsis by Deep Sequencing. Plant Physiol 151, 2120.

45. Hyun, Y., Richter, R., Vincent, C., Martinez-Gallegos, R., Porri, A., and Coupland, G. (2016). Multi-layered Regulation of SPL15 and Cooperation with SOC1 Integrate Endogenous Flowering Pathways at the Arabidopsis Shoot Meristem. Dev Cell 37, 254–266.

46. Journet, E.-P., Neuburger, M., and Douce, R. (1981). Role of Glutamate-oxaloacetate Transaminase and Malate Dehydrogenase in the Regeneration of NAD^+^ for Glycine Oxidation by Spinach leaf Mitochondria. Plant Physiol 67, 467–469.

47. Kadenbach, B., Huttemann, M., Arnold, S., Lee, I., and Bender, E. (2000). Mitochondrial energy metabolism is regulated via nuclear-coded subunits of cytochrome c oxidase. Free Radic Biol Med 29, 211–221.

48. Kim, D., Paggi, J.M., Park, C., Bennett, C., and Salzberg, S.L. (2019). Graph-based genome alignment and genotyping with HISAT2 and HISAT-genotype. Nat Biotechnol 37, 907–915.

49. Kim, S., Mollet, J.-C., Dong, J., Zhang, K., Park, S.-Y., and Lord, E.M. (2003). Chemocyanin, a small basic protein from the lily stigma, induces pollen tube chemotropism. Proc Natl Acad Sci USA 100, 16125.

50. Krämer, U., and Clemens, S. (2005). Functions and homeostasis of zinc, copper, and nickel in plants. In Molecular Biology of Metal Homeostasis and Detoxification, M.J. Tamás and E. Martinoia, eds (Springer Berlin Heidelberg New York), pp. 216–271.

51. Kropat, J., Tottey, S., Birkenbihl, R.P., Depege, N., Huijser, P., and Merchant, S.S. (2005). A regulator of nutritional copper signaling in Chlamydomonas is an SBP domain protein that recognizes the GTAC core of copper response element. Proc Natl Acad Sci USA 102, 18730–18735.

52. Kropat, J., Gallaher, S.D., Urzica, E.I., Nakamoto, S.S., Strenkert, D., Tottey, S., Mason, A.Z., and Merchant, S.S. (2015). Copper economy in Chlamydomonas: prioritized allocation and reallocation of copper to respiration *vs*. photosynthesis. Proc Natl Acad Sci USA 112, 2644–2651.

53. Lambers, H. (1982). Cyanide-resistant respiration: A non-phosphorylating electron transport pathway acting as an energy overflow. Physiol Plant 55, 478–485.

54. Lamesch, P., Berardini, T.Z., Li, D., Swarbreck, D., Wilks, C., Sasidharan, R., Muller, R., Dreher, K., Alexander, D.L., Garcia-Hernandez, M., Karthikeyan, A.S., Lee, C.H., Nelson, W.D., Ploetz, L., Singh, S., Wensel, A., and Huala, E. (2012). The Arabidopsis Information Resource (TAIR): improved gene annotation and new tools. Nucleic Acids Res 40, D1202–1210.

55. Lampropoulos, A., Sutikovic, Z., Wenzl, C., Maegele, I., Lohmann, J.U., and Forner, J. (2013). GreenGate - A Novel, Versatile, and Efficient Cloning System for Plant Transgenesis. PLoS One 8, e83043.

56. Langmead, B., and Salzberg, S.L. (2012). Fast gapped-read alignment with Bowtie 2. Nat Methods 9, 357–359.

57. Langmead, B., Trapnell, C., Pop, M., and Salzberg, S.L. (2009). Ultrafast and memory-efficient alignment of short DNA sequences to the human genome. Genome Biol 10, R25.

58. Lappe, R.R., Baier, J.W., Boehlein, S.K., Huffman, R., Lin, Q., Wattebled, F., Settles, A.M., Hannah, L.C., Borisjuk, L., Rolletschek, H., Stewart, J.D., Scott, M.P., Hennen-Bierwagen, T.A., and Myers, A.M. (2018). Functions of maize genes encoding pyruvate phosphate dikinase in developing endosperm. Proc Natl Acad Sci USA 115, E24–E33.

59. Larronde, F., Krisa, S., Decendit, A., Chèze, C., Deffieux, G., and Mérillon, J.M. (1998). Regulation of polyphenol production in *Vitis vinifera* cell suspension cultures by sugars. Plant Cell Reports 17, 946–950.

60. Lejay, L., Wirth, J., Pervent, M., Cross, J.M., Tillard, P., and Gojon, A. (2008). Oxidative pentose phosphate pathway-dependent sugar sensing as a mechanism for regulation of root ion transporters by photosynthesis. Plant Physiol 146, 2036–2053.

61. Lejay, L., Gansel, X., Cerezo, M., Tillard, P., Muller, C., Krapp, A., von Wiren, N., Daniel-Vedele, F., and Gojon, A. (2003). Regulation of root ion transporters by photosynthesis: functional importance and relation with hexokinase. Plant Cell 15, 2218–2232.

62. Li, L., and Sheen, J. (2016). Dynamic and diverse sugar signaling. Curr Opin Plant Biol 33, 116–125.

63. Li, X., Zhang, H., Ai, Q., Liang, G., and Yu, D. (2016). Two bHLH Transcription Factors, bHLH34 and bHLH104, Regulate Iron Homeostasis in Arabidopsis thaliana. Plant Physiol 170, 2478-2493.

64. Liang, G., Zhang, H., Li, X., Ai, Q., and Yu, D. (2017). bHLH transcription factor bHLH115 regulates iron homeostasis in *Arabidopsis thaliana*. J Exp Bot 68, 1743–1755.

65. Liu, Y., Duan, X., Zhao, X., Ding, W., Wang, Y., and Xiong, Y. (2021). Diverse nitrogen signals activate convergent ROP2-TOR signaling in Arabidopsis. Dev Cell 56, 1283–1295 e1285.

66. Love, M.I., Huber, W., and Anders, S. (2014). Moderated estimation of fold change and dispersion for RNA-seq data with DESeq2. Genome Biol 15, 550.

67. Lunn, J.E., Feil, R., Hendriks, J.H.M., Gibon, Y., Morcuende, R., Osuna, D., Scheible, W.-R., Carillo, P., Hajirezaei, M.-R., and Stitt, M. (2006). Sugar-induced increases in trehalose 6-phosphate are correlated with redox activation of ADPglucose pyrophosphorylase and higher rates of starch synthesis in *Arabidopsis thaliana*. Biochem J 397, 139–148.

68. Machanick, P., and Bailey, T.L. (2011). MEME-ChIP: motif analysis of large DNA datasets. Bioinformatics 27, 1696–1697.

69. Marschner, H., and Marschner, P. (2012). Marschner’s mineral nutrition of higher plants. (London; Waltham, MA: Elsevier/Academic Press).

70. Martin, M. (2011). Cutadapt removes adapter sequences from high-throughput sequencing reads. EMBnet J 17, 3.

71. Moehs, C.P., McElwain, E.F., and Spiker, S. (1988). Chromosomal proteins of *Arabidopsis thaliana*. Pant Mol Biol 11, 507–515.

72. Møller, I.M., Bérczi, A., van der Plas, L.H.W., and Lambers, H. (1988). Measurement of the activity and capacity of the alternative pathway in intact plant tissues: Identification of problems and possible solutions. Physiol Plant 72, 642–649.

73. Nunes, C., Primavesi, L.F., Patel, M.K., Martinez-Barajas, E., Powers, S.J., Sagar, R., Fevereiro, P.S., Davis, B.G., and Paul, M.J. (2013a). Inhibition of SnRK1 by metabolites: Tissue-dependent effects and cooperative inhibition by glucose 1-phosphate in combination with trehalose 6-phosphate. Plant Physiol Biochem 63, 89–98.

74. Nunes, C., O’Hara, L.E., Primavesi, L.F., Delatte, T.L., Schluepmann, H., Somsen, G.W., Silva, A.B., Fevereiro, P.S., Wingler, A., and Paul, M.J. (2013b). The Trehalose 6-Phosphate/SnRK1 Signaling Pathway Primes Growth Recovery following Relief of Sink Limitation. Plant Physiol 162, 1720.

75. O’Leary, B.M., Asao, S., Millar, A.H., and Atkin, Owen K. (2019). Core principles which explain variation in respiration across biological scales. New Phytol 222, 670–686.

76. O’Malley, R.C., Huang, S.C., Song, L., Lewsey, M.G., Bartlett, A., Nery, J.R., Galli, M., Gallavotti, A., and Ecker, J.R. (2016). Cistrome and Epicistrome Features Shape the Regulatory DNA Landscape. Cell 165, 1280–1292.

77. Okonechnikov, K., Conesa, A., and Garcia-Alcalde, F. (2016). Qualimap 2: advanced multi-sample quality control for high-throughput sequencing data. Bioinformatics 32, 292–294.

78. Pietzenuk, B., Markus, C., Gaubert, H., Bagwan, N., Merotto, A., Bucher, E., and Pecinka, A. (2016). Recurrent evolution of heat-responsiveness in Brassicaceae COPIA elements. Genome Biol 17, 209.

79. Ponnu, J., Schlereth, A., Zacharaki, V., Dzialo, M.A., Abel, C., Feil, R., Schmid, M., and Wahl, V. (2020). The trehalose 6-phosphate pathway impacts vegetative phase change in *Arabidopsis thaliana*. Plant J 104, 768–780.

80. Quinn, J.M., and Merchant, S.S. (1995). Two copper-responsive elements associated with the Chlamydomonas *Cyc6* gene function as targets for transcriptional activators. Plant Cell 7, 623–628.

81. R Core Team. (2019). R: A language and environment for statistical computing. R Foundation for Statistical Computing, Vienna, Austria.

82. Rabino, I., and Mancinelli, A.L. (1986). Light, Temperature, and Anthocyanin Production. Plant Physiol 81, 922–924.

83. Rae, T.D., Schmidt, P.J., Pufahl, R.A., Culotta, V.C., and O’Halloran, T.V. (1999). Undetectable intracellular free copper: the requirement of a copper chaperone for superoxide dismutase. Science 284, 805–808.

84. Rahmati Ishka, M., and Vatamaniuk, O.K. (2020). Copper deficiency alters shoot architecture and reduces fertility of both gynoecium and androecium in Arabidopsis thaliana. Plant Direct 4, e00288.

85. Ravet, K., Danford, F.L., Dihle, A., Pittarello, M., and Pilon, M. (2011). Spatiotemporal analysis of copper homeostasis in *Populus trichocarpa* reveals an integrated molecular remodeling for a preferential allocation of copper to plastocyanin in the chloroplasts of developing leaves. Plant Physiol 157, 1300–1312.

86. Redinbo, M.R., Yeates, T.O., and Merchant, S. (1994). Plastocyanin: structural and functional analysis. J Bioenerg Biomembr 26, 49–66.

87. Reimann, C., Birke, M., Demetriades, A., Filzmoser, P., and O’Connor, P. (2014). Chemistry of Europe’s agricultural soils - Part A: Methodology and interpretation of the GEMAS data set. (Hannover: Schweizerbarth).

88. Ren, L., and Tang, G. (2012). Identification of sucrose-responsive microRNAs reveals sucrose-regulated copper accumulations in an SPL7-dependent and independent manner in *Arabidopsis thaliana*. Plant Sci 187, 59–68.

89. Robinson, N.J., and Winge, D.R. (2010). Copper metallochaperones. Annu Rev Biochem 79, 537–562.

90. Roman, A., Li, X., Deng, D., Davey, J.W., James, S., Graham, I.A., and Haydon, M.J. (2021). Superoxide is promoted by sucrose and affects amplitude of circadian rhythms in the evening. Proc Natl Acad Sci USA 118.

91. Romera-Branchat, M., Severing, E., Pocard, C., Ohr, H., Vincent, C., Nee, G., Martinez-Gallegos, R., Jang, S., Andres, F., Madrigal, P., and Coupland, G. (2020). Functional Divergence of the Arabidopsis Florigen-Interacting bZIP Transcription Factors FD and FDP. Cell Rep 32, 107966.

92. Schluepmann, H., Pellny, T., van Dijken, A., Smeekens, S., and Paul, M. (2003). Trehalose 6-phosphate is indispensable for carbohydrate utilization and growth in *Arabidopsis thaliana*. Proc Natl Acad Sci USA 100, 6849–6854.

93. Schubert, M., Petersson, U.A., Haas, B.J., Funk, C., Schroder, W.P., and Kieselbach, T. (2002). Proteome map of the chloroplast lumen of *Arabidopsis thaliana*. J Biol Chem 277, 8354–8365.

94. Schulten, A., Bytomski, L., Quintana, J., Bernal, M., and Kramer, U. (2019). Do Arabidopsis Squamosa promoter binding Protein-Like genes act together in plant acclimation to copper or zinc deficiency? Plant Direct 3, e00150.

95. Schwarz, S., Grande, A.V., Bujdoso, N., Saedler, H., and Huijser, P. (2008). The microRNA regulated SBP-box genes *SPL9* and *SPL15* control shoot maturation in Arabidopsis. Plant Mol Biol 67, 183–195.

96. Shen, W., Wei, Y., Dauk, M., Tan, Y., Taylor, D.C., Selvaraj, G., and Zou, J. (2006). Involvement of a Glycerol-3-Phosphate Dehydrogenase in Modulating the NADH/NAD^+^ Ratio Provides Evidence of a Mitochondrial Glycerol-3-Phosphate Shuttle in Arabidopsis. Plant Cell 18, 422–441.

97. Shikanai, T., Müller-Moulé, P., Munekage, Y., Niyogi, K.K., and Pilon, M. (2003). PAA1, a P-type ATPase of Arabidopsis, functions in copper transport in chloroplasts. Plant Cell 15, 1333–1346.

98. Sinclair, S.A., Larue, C., Bonk, L., Khan, A., Castillo-Michel, H., Stein, R.J., Grolimund, D., Begerow, D., Neumann, U., Haydon, M.J., and Krämer, U. (2017). Etiolated Seedling Development Requires Repression of Photomorphogenesis by a Small Cell-Wall-Derived Dark Signal. Curr Biol 27, 3403–3418 e3407.

99. Solfanelli, C., Poggi, A., Loreti, E., Alpi, A., and Perata, P. (2006). Sucrose-specific induction of the anthocyanin biosynthetic pathway in Arabidopsis. Plant Physiol 140, 637–646.

100. Sommer, F., Kropat, J., Malasarn, D., Grossoehme, N.E., Chen, X., Giedroc, D.P., and Merchant, S.S. (2010). The CRR1 nutritional copper sensor in Chlamydomonas contains two distinct metal-responsive domains. Plant Cell 22, 4098–4113.

101. Stief, A., Altmann, S., Hoffmann, K., Pant, B.D., Scheible, W.-R., and Bäurle, I. (2014). Arabidopsis miR156 Regulates Tolerance to Recurring Environmental Stress through SPL Transcription Factors. Plant Cell 26, 1792.

102. Stone, J.M., Liang, X., Nekl, E.R., and Stiers, J.J. (2005). Arabidopsis AtSPL14, a plant-specific SBP-domain transcription factor, participates in plant development and sensitivity to fumonisin B1. Plant J 41, 744–754.

103. Storey, J.D., Bass, A.J., Dabney, A., and Robinson, D. (2019). qvalue: Q-value estimation for false discovery rate control. R package version 2.14.1.

104. Teng, S., Keurentjes, J., Bentsink, L., Koornneef, M., and Smeekens, S. (2005). Sucrose-specific induction of anthocyanin biosynthesis in Arabidopsis requires the *MYB75/PAP1* gene. Plant Physiol 139, 1840–1852.

105. Towbin, H., Staehelin, T., and Gordon, J. (1979). Electrophoretic transfer of proteins from polyacrylamide gels to nitrocellulose sheets: procedure and some applications. Proc Natl Acad Sci USA 76, 4350–4354.

106. Unte, U.S., Sorensen, A.-M., Pesaresi, P., Gandikota, M., Leister, D., Saedler, H., and Huijser, P. (2003). *SPL8*, an SBP-box gene that affects pollen sac development in Arabidopsis. Plant Cell 15, 1009–1019.

107. Urano, D., Phan, N., Jones, J.C., Yang, J., Huang, J., Grigston, J., Taylor, J.P., and Jones, A.M. (2012). Endocytosis of the seven-transmembrane RGS1 protein activates G-protein-coupled signalling in Arabidopsis. Nat Cell Biol 14, 1079–1088.

108. Varkonyi-Gasic, E., Wu, R., Wood, M., Walton, E.F., and Hellens, R.P. (2007). Protocol: a highly sensitive RT-PCR method for detection and quantification of microRNAs. Plant Methods 3, 12.

109. Wahl, V., Ponnu, J., Schlereth, A., Arrivault, S., Langenecker, T., Franke, A., Feil, R., Lunn, J.E., Stitt, M., and Schmid, M. (2013). Regulation of flowering by trehalose-6-phosphate signaling in *Arabidopsis thaliana*. Science 339, 704–707.

110. Wang, H.-Y., Klatte, M., Jakoby, M., Bäumlein, H., Weisshaar, B., and Bauer, P. (2007). Iron deficiency-mediated stress regulation of four subgroup Ib BHLH genes in *Arabidopsis thaliana*. Planta 226, 897–908.

111. Wang, J.-W., Schwab, R., Czech, B., Mica, E., and Weigel, D. (2008). Dual Effects of miR156-Targeted SPL Genes and CYP78A5/KLUH on Plastochron Length and Organ Size in *Arabidopsis thaliana*. Plant Cell 20, 1231.

112. Wang, J., Zhou, L., Shi, H., Chern, M., Yu, H., Yi, H., He, M., Yin, J., Zhu, X., Li, Y., Li, W., Liu, J., Wang, J., Chen, X., Qing, H., Wang, Y., Liu, G., Wang, W., Li, P., Wu, X., Zhu, L., Zhou, J.-M., Ronald, P.C., Li, S., Li, J., and Chen, X. (2018). A single transcription factor promotes both yield and immunity in rice. Science 361, 1026.

113. Wang, J.W., Czech, B., and Weigel, D. (2009). miR156-regulated SPL transcription factors define an endogenous flowering pathway in *Arabidopsis thaliana*. Cell 138, 738–749.

114. Weigel, M., Varotto, C., Pesaresi, P., Finazzi, G., Rappaport, F., Salamini, F., and Leister, D. (2003). Plastocyanin is indispensable for photosynthetic electron flow in *Arabidopsis thaliana*. J Biol Chem 278, 31286–31289.

115. Weiss, D. (2000). Regulation of flower pigmentation and growth: Multiple signaling pathways control anthocyanin synthesis in expanding petals. Physiol Plant 110, 152–157.

116. Woeste, K.E., and Kieber, J.J. (2000). A strong loss-of-function mutation in RAN1 results in constitutive activation of the ethylene response pathway as well as a rosette-lethal phenotype. Plant Cell 12, 443–455.

117. Wu, G., and Poethig, R.S. (2006). Temporal regulation of shoot development in *Arabidopsis thaliana* by miR156 and its target *SPL3*. Development 133, 3539–3547.

118. Wu, G., Park, M.Y., Conway, S.R., Wang, J.W., Weigel, D., and Poethig, R.S. (2009). The sequential action of miR156 and miR172 regulates developmental timing in Arabidopsis. Cell 138, 750–759.

119. Xing, S., Salinas, M., Hohmann, S., Berndtgen, R., and Huijser, P. (2010). miR156-targeted and nontargeted SBP-box transcription factors act in concert to secure male fertility in Arabidopsis. Plant Cell 22, 3935–3950.

120. Xing, S., Salinas, M., Garcia-Molina, A., Hohmann, S., Berndtgen, R., and Huijser, P. (2013). *SPL8* and miR156-targeted *SPL genes* redundantly regulate Arabidopsis gynoecium differential patterning. Plant J 75, 566–577.

121. Xiong, Y., and Sheen, J. (2012). Rapamycin and glucose-target of rapamycin (TOR) protein signaling in plants. J Biol Chem 287, 2836–2842.

122. Xiong, Y., McCormack, M., Li, L., Hall, Q., Xiang, C., and Sheen, J. (2013). Glucose-TOR signalling reprograms the transcriptome and activates meristems. Nature 496, 181–186.

123. Xu, M., Hu, T., Zhao, J., Park, M.Y., Earley, K.W., Wu, G., Yang, L., and Poethig, R.S. (2016). Developmental Functions of miR156-Regulated *SQUAMOSA PROMOTER BINDING PROTEIN-LIKE (SPL)* Genes in *Arabidopsis thaliana*. PLoS Genet 12, e1006263.

124. Yamaguchi, A., Wu, M.F., Yang, L., Wu, G., Poethig, R.S., and Wagner, D. (2009). The microRNA-regulated SBP-Box transcription factor SPL3 is a direct upstream activator of LEAFY, FRUITFULL, and APETALA1. Dev Cell 17, 268-278.

125. Yamaguchi, N., Winter, C.M., Wu, M.-F., Kwon, C.S., William, D.A., and Wagner, D. (2014a). PROTOCOLS: Chromatin Immunoprecipitation from Arabidopsis Tissues. The Arabidopsis Book 12, e0170–e0170.

126. Yamaguchi, N., Winter, C.M., Wu, M.F., Kanno, Y., Yamaguchi, A., Seo, M., and Wagner, D. (2014b). Gibberellin acts positively then negatively to control onset of flower formation in Arabidopsis. Science 344, 638–641.

127. Yamasaki, H., Hayashi, M., Fukazawa, M., Kobayashi, Y., and Shikanai, T. (2009). SQUAMOSA Promoter Binding Protein-Like7 Is a Central Regulator for Copper Homeostasis in Arabidopsis. Plant Cell 21, 347–361.

128. Yamasaki, H., Abdel-Ghany, S.E., Cohu, C.M., Kobayashi, Y., Shikanai, T., and Pilon, M. (2007). Regulation of copper homeostasis by micro-RNA in Arabidopsis. J Biol Chem 282, 16369–16378.

129. Yan, J., Chia, J.-C., Sheng, H., Jung, H.-I., Zavodna, T.-O., Zhang, L., Huang, R., Jiao, C., Craft, E.J., Fei, Z., Kochian, L.V., and Vatamaniuk, O.K. (2017). Arabidopsis Pollen Fertility Requires the Transcription Factors CITF1 and SPL7 That Regulate Copper Delivery to Anthers and Jasmonic Acid Synthesis. Plant Cell 29, 3012–3029.

130. Yang, L., Conway, S.R., and Poethig, R.S. (2011). Vegetative phase change is mediated by a leaf-derived signal that represses the transcription of miR156. Development 138, 245–249.

131. Yang, L., Xu, M., Koo, Y., He, J., and Poethig, R.S. (2013). Sugar promotes vegetative phase change in *Arabidopsis thaliana* by repressing the expression of *MIR156A* and *MIR156C*. eLife 2, e00260.

132. Yruela, I. (2013). Transition metals in plant photosynthesis. Metallomics 5, 1090–1109.

133. Yu, N., Cai, W.-J., Wang, S., Shan, C.-M., Wang, L.-J., and Chen, X.-Y. (2010). Temporal Control of Trichome Distribution by MicroRNA156-Targeted *SPL* Genes in *Arabidopsis thaliana*. Plant Cell 22, 2322.

134. Yu, S., Cao, L., Zhou, C.-M., Zhang, T.-Q., Lian, H., Sun, Y., Wu, J., Huang, J., Wang, G., and Wang, J.-W. (2013). Sugar is an endogenous cue for juvenile-to-adult phase transition in plants. eLife 2, e00269.

135. Zhai, Z., Keereetaweep, J., Liu, H., Feil, R., Lunn, J.E., and Shanklin, J. (2018). Trehalose 6-Phosphate Positively Regulates Fatty Acid Synthesis by Stabilizing WRINKLED1. Plant Cell 30, 2616–2627.

136. Zhang, H., and Li, L. (2013). SQUAMOSA promoter binding protein-like7 regulated microRNA408 is required for vegetative development in Arabidopsis. Plant J 74, 98–109.

137. Zhang, H., Zhao, X., Li, J., Cai, H., Deng, X.W., and Li, L. (2014). MicroRNA408 is critical for the HY5-SPL7 gene network that mediates the coordinated response to light and copper. Plant Cell 26, 4933–4953.

138. Zhang, Y., Primavesi, L.F., Jhurreea, D., Andralojc, P.J., Mitchell, R.A.C., Powers, S.J., Schluepmann, H., Delatte, T., Wingler, A., and Paul, M.J. (2009). Inhibition of SNF1-Related Protein Kinase1 Activity and Regulation of Metabolic Pathways by Trehalose-6-Phosphate. Plant Physiol 149, 1860–1871.

139. Zhang, Y., Liu, T., Meyer, C.A., Eeckhoute, J., Johnson, D.S., Bernstein, B.E., Nusbaum, C., Myers, R.M., Brown, M., Li, W., and Liu, X.S. (2008). Model-based Analysis of ChIP-Seq (MACS). Genome Biol 9, R137.

